# Simulation Enabled Search for Explanatory Mechanisms of the Fracture Healing Process

**DOI:** 10.1101/216523

**Authors:** Ryan C. Kennedy, Meir Marmor, Ralph Marcucio, C. Anthony Hunt

**Affiliations:** Department of Bioengineering and Therapeutic Sciences, University of California, San Francisco, CA 94143; Department of Orthopaedic Surgery, San Francisco General Hospital Orthopaedic Trauma Institute, University of California, San Francisco, CA 94110

## Abstract

A significant portion of bone fractures fail to heal properly, increasing healthcare costs. Advances in fracture management have slowed because translational barriers have limited generation of mechanism-based explanations for the healing process. When uncertainties are numerous, analogical modeling can be an effective strategy for developing plausible explanations of complex phenomena. We demonstrate the feasibility of engineering analogical models in software to provide plausible biomimetic explanations for how fracture healing may progress. Concrete analogical models – Callus Analogs – were created using the MASON simulation toolkit. We designated a Target Region initial state within a characteristic tissue section of mouse tibia fracture at day-7 and posited a corresponding day-10 Target Region final state. The goal was to discover a coarse-grain analog mechanism that would enable the discretized initial state to transform itself into the corresponding Target Region final state, thereby providing a new way to study the healing process. One of nine quasi-autonomous Tissue Unit types is assigned to each grid space, which maps to an 80×80 µm region of the tissue section. All Tissue Units have an opportunity each time step to act based on individualized logic, probabilities, and information about adjacent neighbors. Action causes transition from one Tissue Unit type to another, and simulation through several thousand time steps generates a coarse-grain analog – a theory – of the healing process. We prespecified a minimum measure of success: simulated and actual Target Region states achieve ≥ 70% Similarity. We used an iterative protocol to explore many combinations of Tissue Unit logic and action constraints. Workflows progressed through four stages of analog mechanisms. Similarities of 73-90% were achieved for Mechanisms 2-4. The range of Upper-Level similarities increased to 83-94% when we allowed for uncertainty about two Tissue Unit designations. We have demonstrated how Callus Analog experiments provide domain experts with a new medium and tools for thinking about and understanding the fracture healing process.

**Author summary:** Translational barriers have limited the generation of mechanism-based explanations of fracture healing processes. Those barriers help explain why, to date, biological therapeutics have had only a minor impact on fracture management. New approaches are needed, and we present one that is intended to overcome those barriers incrementally. We created virtual Callus Analogs to simulate how the histologic appearance of a mouse fracture callus may transition from day-7 to day-10. Callus Analogs use software-based model mechanisms. Simulation experiments enable challenging and improving those model mechanisms. During execution, model mechanism operation provides a coarse-grain explanation (a theory) of a four-day portion of the healing process. Simulated day-10 callus histologic images achieved 73-94% similarity to a corresponding day-10 fracture callus image, thus demonstrating feasibility. Simulated healing provides a new perspective on the actual healing process and a new way of thinking about plausible fracture-healing processes. Our working hypothesis is that the approach can be extended to cover more of the healing process while making features of simulated and actual fracture healing increasingly analogous. The methods presented are intended to be extensible to other research areas that use histologic analysis to investigate and explain tissue level phenomena.

## Introduction

Annually, there are approximately 15 million fractures in the United States, and a significant portion (10-15%) fail to heal properly [1]. Both numbers and costs are predicted to increase as the population ages and the number of osteoporosis-related fractures increases [2]. Therefore, developing intervention strategies to stimulate fracture healing is expected to positively impact health. Many of the advances made in fracture management in recent years were in mechanical stabilization and biologic bone augmentation materials such as autogenous bone graft, synthetic bone ceramics, or demineralized bone matrix [3]. The clinical impact of biological therapeutic agents, such as bone morphogenetic proteins, has fallen short of expectations for largely unknown reasons [4]. It is noteworthy that the gold standard, and most commonly used strategy for fracture nonunion treatment, autogenous bone graft, has not changed in the last 100 years [3, 5]. Introductions of new therapeutics have slowed despite expanded research [6]. Such ineffectiveness reflects significant translation barriers. The problem is not unique to fracture-healing research; it is encountered within many research domains [7, 8]. A translation barrier is created when knowledge and explanations of complex medical processes, such as fracture healing, are insufficient to identify a reliable, efficacious intervention strategy. A goal of the research described herein is to develop a new simulation-based approach for improving mechanism-oriented explanations of the fracture healing process.

Fracture healing is described as comprising two phases and three stages that overlap temporally over several weeks: anabolic and catabolic phases; and inflammatory, endochondral, and coupled remodeling stages. The dominant cell types and subprocesses [9] change as healing progresses. Recent analyses of transcriptomes present during fracture healing have shown that most of the genes and signaling pathways that are involved in skeletal development in embryos are also expressed in cells of the fractured callus [10]. Consequently, some pathway components have become the focus of empirical research efforts to develop therapeutic interventions [9], despite the fact that there is no explanatory model of the fracture healing process.

Core phenomena of embryogenesis and some types of tissue regeneration include the evolving small- and large-scale patterns that are readily apparent in recorded images. There has been considerable progress in developing goal-directed mechanism-oriented explanations for those phenomena [11]. However, stained tissue sections of mouse tibia fractures obtained at intervals of several days lack the hallmarks of orderly, organized evolving phenomena exhibited by embryogenesis. The strikingly less organized callus tissue obscures the ongoing order of the various subprocesses and their mechanisms. Part of the problem traces to limitations of experimentation. Healing of mouse tibia fractures typically spans four-to-five weeks. A major complication is that, within the same experiment, no two fractures are the same. Although the healing phenomenon is the same, the healing subprocesses within each callus as they unfold are unique. Large observational gaps coupled with the necessary limitations of standard histological techniques means that informative processes or phenomena may be missed. It is also plausible that informative phenomena—patterns and features—are being observed and recorded, but are not yet recognized as such.

Analogous circumstances have existed in non-biological domains, and significant progress has been achieved by using computational and grid-based simulation methods to provide plausible representations of the missing processes and phenomena. For example, looking for improved insight into processes occurring at the interface of ecology and geomorphology, Fonstad opined, “we have thousands of such images, but no theories in geomorphology nor ecology can fully explain the patterns in any of them” [12].

The fact that callus mechanisms have been successfully healing bone fractures for more than 150 million years [13] implies the existence of a well-orchestrated, robust process. Similarities of callus and embryonic transcriptomes support that inference [2]. If we accept the premise that fracture healing is a well-orchestrated, robust process, then we need to answer this question: how can we begin developing a theory about the healing process—even if initially coarse and somewhat abstract—so that we can begin theorizing about its orchestration? A clearly described phenomenon is a precondition for developing a theory intended to explain that phenomenon (see S1 File). However, we do not yet have a clear temporal description of the fracture healing process, or even for portions of the process. We do, however, have detailed descriptions of features of the process at different stages.

With current technology, it is not feasible to measure callus development continuously. Likewise, it is infeasible to track the changing variety of local structures and cell types. Must we plead for more data, and then come back to the problem in another decade or two? Absent a plausible explanation and theory to test, more data may not be the answer. In discussing comparable issues at the ecology-geomorphology interface, Fonstad observed that, “both of these disciplines are data-rich … it is immediately apparent that both of these disciplines are far more theory-poor” [12]. Fracture-healing research is handicapped because it is relatively data-poor *and* theory-poor. So, although we can draw inspiration from the explanatory simulation methods used by Fonstad and others, their models and simulation methods are not directly applicable in advancing fracture-healing research.

Given the increasing interest in increasing the clinical relevance of modeling and simulation research, it is not surprising that the number of such reports in which authors utilize histology images to support face validation and/or guide calibrations is increasing. The following are three recent examples. Marino et al. [14] utilized their model of lung granuloma formation to compare in silico granulomas to those of the nonhuman primate *Macaca fascicularis*. Gardiner et al. [15] utilized an agent-oriented particle system at various granularities to simulate mechanical behaviors of cells and tissues. Simulations using selected parameterizations bore a remarkable resemblance to histological observations of an epithelial layer, cell clusters, and single cells. Ziraldo et al. describe an agent-based model of ischemia/reperfusion-induced inflammation coupled with pressure ulcer formation and progression in humans suffering from a spinal cord injury [16]. Serial photographic images spanning several clinical stages were used to calibrate progression and healing of virtual pressure ulcers. Virtual pressure ulcers were interrogated to explore how and when an irritation might resolve or become chronic.

The prospect of pulling together a start-to-finish tissue-level mechanism-oriented description of a fracture healing process, even one that is initially coarse-grain, seems distant. Why? It is a consequence of four interrelated obstacles arising from fracture-healing research using rodent models.

1. Missing system information looms largest. Knowledge about the healing process is still sparse compared to that available in other areas of research, such as embryonic skeletal development. Histological observations made during even the most thorough wet-lab experiments cover only a small fraction of the temporal space of callus behaviors. Current wet-lab methods provide snapshots, spaced days apart, of the dynamic, evolving, and multifaceted healing process.
2. Because each callus is unique, there is considerable inter-callus variability. Yet the healing process is sufficiently robust so that callus variability is easily accommodated: absent disruption, bone restoration is always the end state. Quantitative models of sequential features are needed to begin developing an overall explanatory theory. However, those features may not be easily identified when a higher-level process comprises many subprocesses.
3. Informative data that characterizes sequential feature changes are sparse at best. Most of the available data are measurements made at cell and intracellular levels.
4. Pervasive uncertainties beyond callus variability obscure process order. Each mouse fracture can be observed histologically only once. In addition, the pace of healing processes is subject to several biological factors, such as age [17], along with a number of external factors [9], notably diabetes [18], smoking [19], and vitamin D deficiency [20]. Controlling such factors both within and between experiments can be challenging and expensive. Consequently, distinguishing causes from effects can be problematic.

Given those obstacles, knowledge is insufficient to describe, much less begin building a conventional molecular and cellular biology-based model of fracture healing. The most pressing current need is to develop strategies and methods to workaround and eventually overcome each of the above four obstacles. We conjectured that the software-based model mechanism methods, which we have used successfully in other contexts (e.g., see [21–26]), could provide the foundation for such strategies, even though, for those earlier applications, considerably more mechanism related fine grain knowledge was available. Briefly stated, the cited software-based model mechanism approach begins with a target phenomenon. We build an extant (actually existing, observable), working mechanism in software that is parsimonious and, based on a similarity criterion, exhibits essentially the same phenomenon. Doing so requires making no assumptions about the biology.

But even when the mechanism is kept coarse grain, the space of possible software mechanisms capable of generating essentially the same phenomenon can be huge. So, biologically inspired requirements and constraints along with mechanism granularity limits are imposed incrementally to shrink and constrain possible model mechanism space. That process shrinks a large set of possible coarse grain mechanisms into a much smaller set of plausible, incrementally more likely and increasingly biomimetic, model mechanisms.

For fracture healing we envision simulations generating plausible scenarios for how discretized features of a callus tissue section on one day might progressively transform itself into the corresponding tissue section features—target features—observed several days later. Wet-lab experiments can target differences in two mechanisms, where the resulting new evidence is expected to support one mechanism and falsify the other (as in [27]), further shrinking plausible mechanism space. At that stage, the surviving software mechanism can stand as a coarse grain theory for how a portion of mouse tibia fracture healing is occurring.

To begin eroding the four obstacles in meaningful ways requires coupling the preceding methods with an important new capability: use image interpolation strategies to build plausible sequential image models of the same fracture at different stages of the healing process. A requisite for an interpolation strategy is having and aligning discretized coarse-grain models of tissue section images of tibia fractures from different mice at different times.

We report results of a focused demonstration that meets the above requirements. We present results of workflows that support the feasibility of the approach, while also bringing its weaknesses into focus. For this demonstration, we limited attention to the critical interval from day-7 to day-10 during healing of a mouse tibia fracture and focused on discretized models of specific tissue sections on both days. From the latter, we obtained the initial state and final target state for our simulations. Biomimetic software mechanisms involving actions of quasi-autonomous tissue units spanning, typically, 5,000-6,000 time steps are responsible for simulated healing. Similarities (defined in Methods) between simulated and referent final states ranged from > 70% to > 93%, depending on the stringency of the similarity criterion. Despite the narrow focus, it is clear that a major benefit of the approach is demonstrating that simulation experiments can enable discovering, challenging, and improving theories of healing subprocesses.

Because our approach and methods are unconventional, somewhat new, and still evolving, we present that information next under Methods to provide the context needed to present and discuss results. There are weaknesses and limitations associated with every aspect of our approach. Some are identified in Methods, and others are addressed under Discussion. We undertook this demonstration with the expectation that the more successful methods could be repurposed to begin lowering similar barriers faced within some domains of disease progression research.

## Methods

### Approach

We begin with a synopsis of our approach from a workflow perspective, as diagrammed in Fig 1. We then provide details on methods used during each of the six stages. In several places, we also provide essential background information that influenced decisions for a particular stage. Words, such as tissue, mechanism, healing, and process, are used in discussing actual mouse tibia fracture healing and corresponding simulations. To reduce confusion, we capitalize those words hereafter when discussing Callus Analogs.

**Fig 1.**
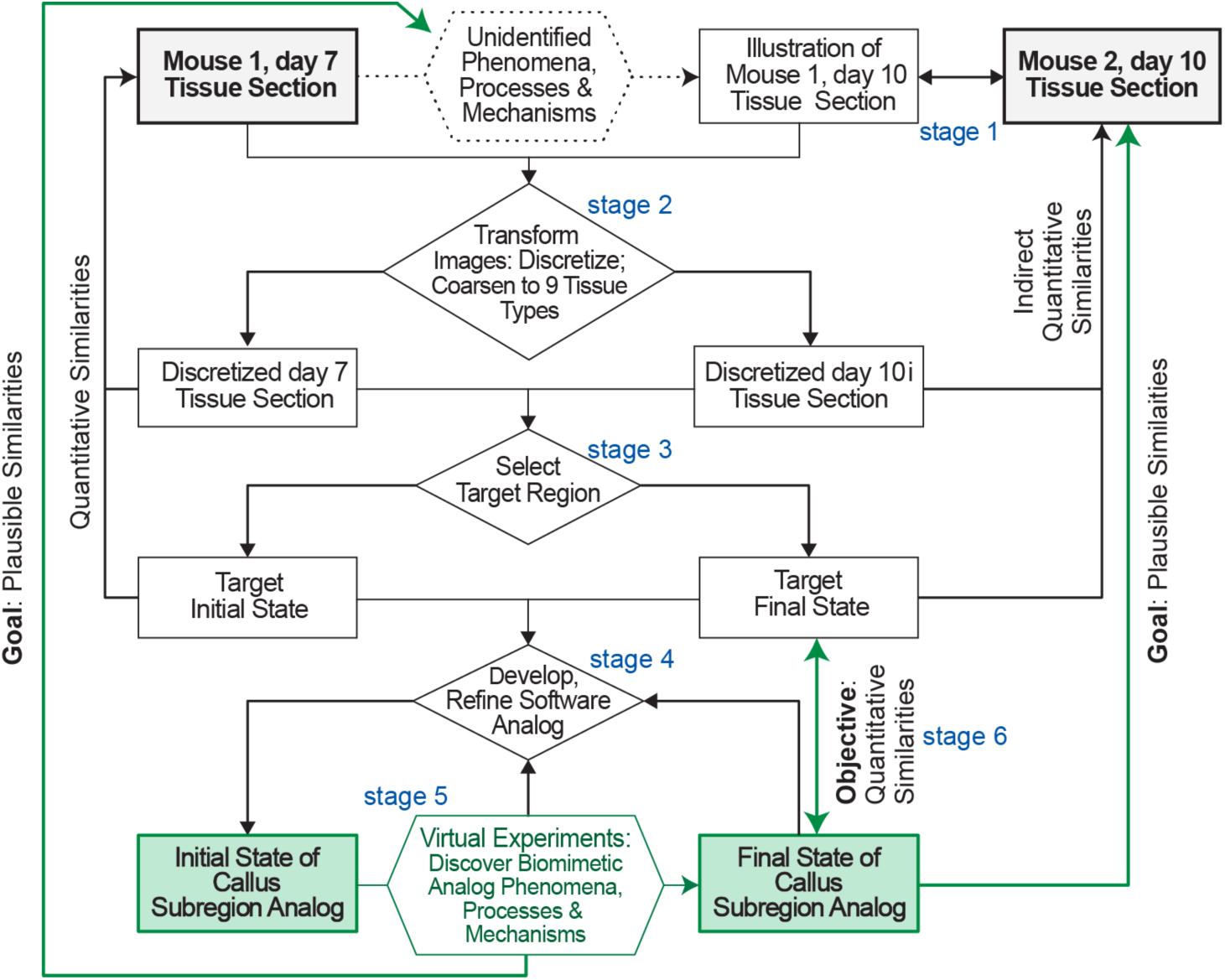
This workflow sketch identifies key features and stages of our approach. We have two concurrent long-term goals. 1) Build the case for plausible similarities—analogies—between Callus Subregion Analog final states (green boxes) and stained tissue sections in Fig 2 obtained from mouse 2 after 10 days of healing (shaded gray boxes). 2) Also build the case that strong analogies can exist between Callus Analog Mechanisms and Processes occurring during execution and corresponding mouse callus healing mechanisms and processes at comparable granularities. The objective of this work is stage 6: demonstrate quantitative similarities between simulated final states and the day-10i Target Region.

We focus on the day-7 to day-10 interval of mouse tibia fracture healing because histomorphological evidence indicates that the relative contributions of chondrogenesis and osteogenesis may undergo important changes during that interval. The goal is to develop a concrete, quantitative (and thus challengeable) but partially coarse-grain theory that explains how characteristic tissue level features on day-7 are being transformed into corresponding features observed on day-10. The discovery effort would be greatly simplified if we could obtain collocated day-7 and day-10 tissue sections from the same callus (mouse 1), but that is infeasible. Instead, we used an evidence-based illustration (created by coauthor M.M.) of an envisioned tissue section of the mouse 1 callus on day-10 at the same callus location as the day-7 tissue section. Three domain experts (see Acknowledgements) judged it plausible and acceptable. Hereafter, we refer to the illustration as the day-10i tissue section. A square grid was used to discretize the day-7 and day-10i images. The area of tissue at each grid location was labeled as one of nine tissue types, based on staining and preponderance of cell types within that space. The result (stage 2) was a discretized coarse-grain model of the day-7 and day-10i tissue sections.

Because we are at the beginning of this explanatory discovery process, we needed and selected a target region on which to focus (discussed further below under Target Region). From a simulation perspective, the target region has an initial state, which maps to the day-7 tissue section, and a corresponding final state, which maps to day-10i tissue section. During stage 4 we used the MASON simulation toolkit [28] to create a 2D 25×25 Target Region initial state, in which objects representing tissue units are assigned to each grid space. We start with a 2D Target Region to limit uncertainty in tissue type identification and to adhere to our parsimony guideline. Stage 5 efforts focused on answering the following question: how do we enable the Target Region initial state to transform itself so that the arrangement of tissue types mimics the Target Region final state? The steps followed to answer that question involved iterative refinements and had two objectives. 1) Explore logic to be used by simulated tissue units that enable them to successfully transition into biomimetic final states. 2) In doing so, keep the logic simple and avoid process features that may appear non-biomimetic. Once we had evidence that reasonably biomimetic final states were achievable, we shifted attention to improving the simulated healing process sufficiently to achieve the following quantitative target Similarity value (stage 6): compositional and organizational similarity between simulated and Target Region final state is ≥ 70%. So doing would support the feasibility of achieving the long-term Fig 1 goals. A simulation that uses concrete objects (simulated tissue units) to generate a process that is analogous to callus healing in several ways. It is a software analog of the healing process. We call the parameterized software a Callus Subregion Analog. Hereafter, for convenience, we use Callus Analog and in some places simply Analog.

### Tissue section images and their interpolation

Histologic slides of sagittal sections through mouse tibia calluses at various stages of healing were available from a previous study. The sections were stained using Hall-Brunt Quadruple to highlight tissue, bone, and cartilage. Shown in Fig 2 are the tissue sections from mouse 1 on day-7, and from mouse 2 on day-10. Coauthor R.M. selected them because they have similar fracture features and exhibit all characteristic callus features.

**Fig 2.**
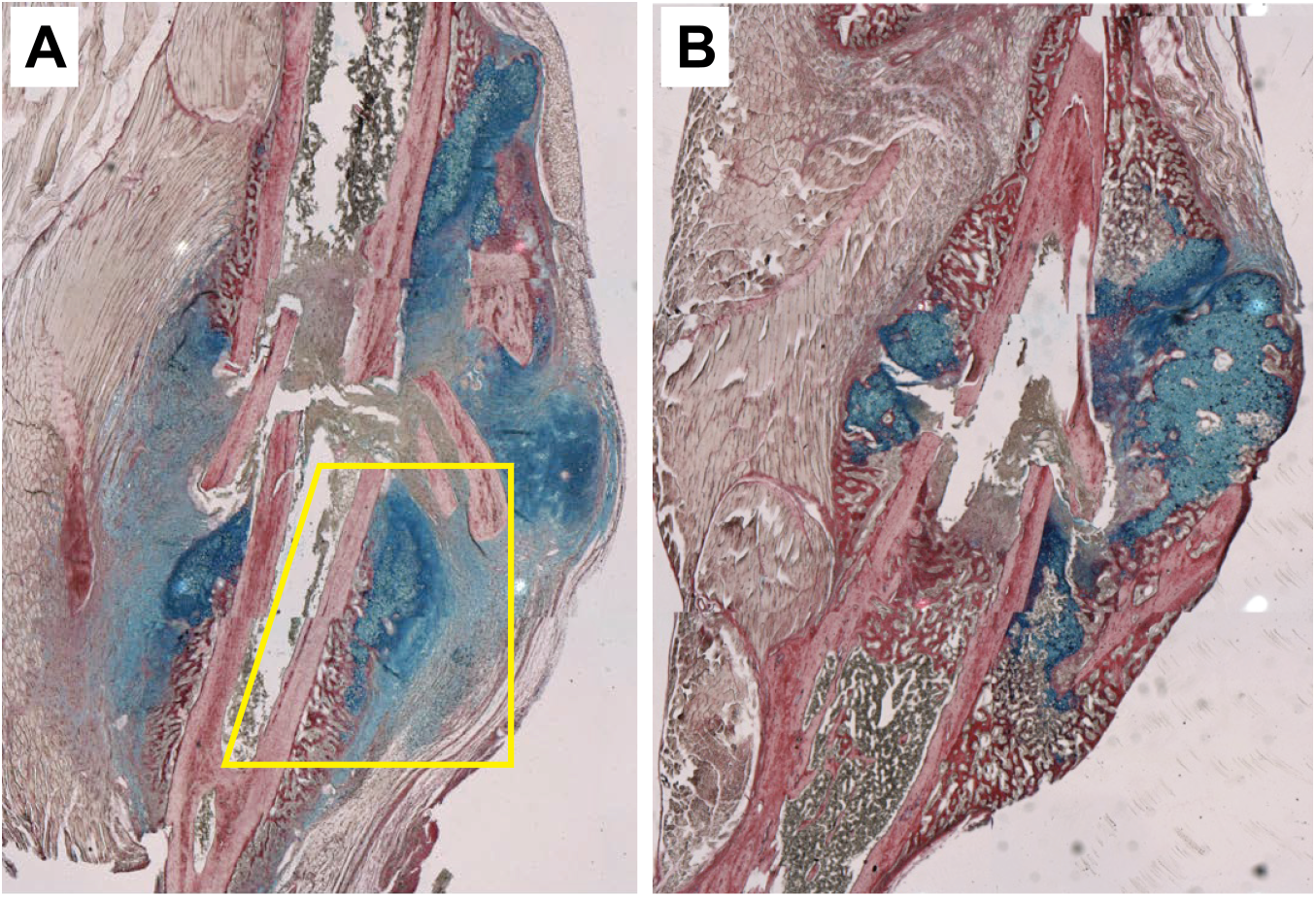
Shown are images of sagittal sections through mouse tibia calluses that were stained for tissue, bone, and cartilage using Hall-Brunt Quadruple. (A) Mouse 1 on day-7; we decided to locate the Target Region within yellow-boxed area. (B) Mouse 2 on day-10.

The following nine distinct microscopic tissue types are common to all normal mouse tibia calluses, beginning before day-7 and extending beyond day-10. We assigned a different color to each tissue type, which was used them to colorized a discretized version of Fig 2A.

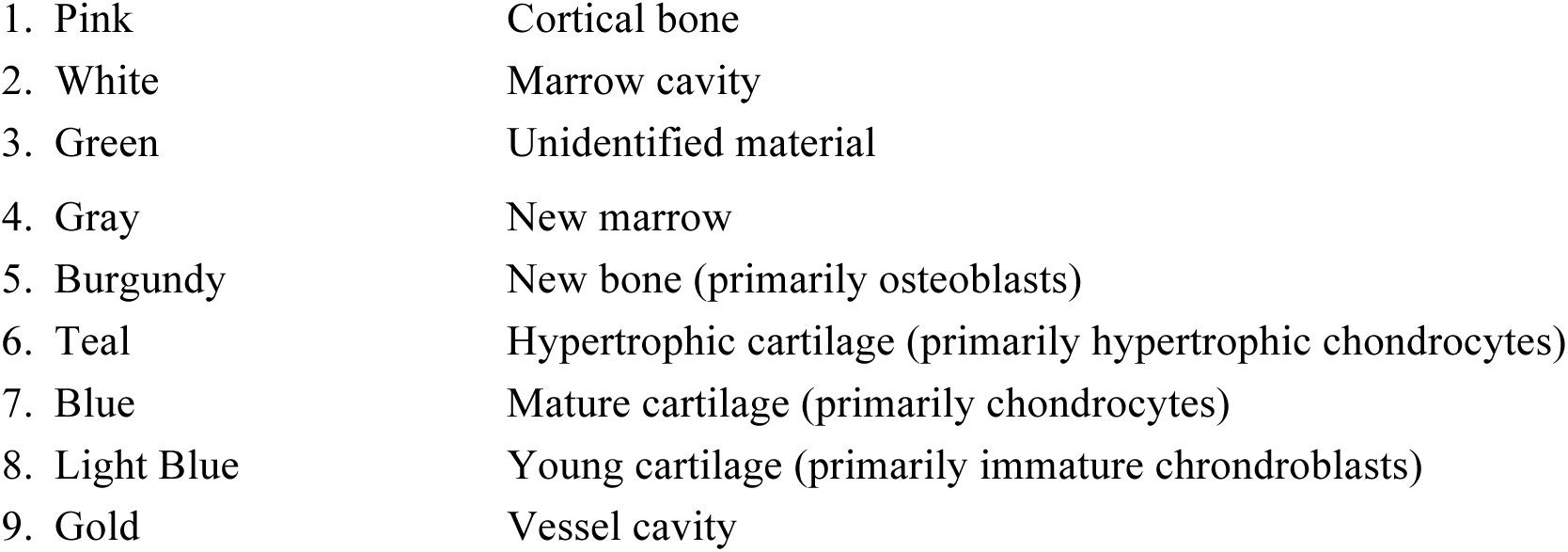

A microscopic area of callus containing about 20 or more cells can be distinguished as being either new marrow (4), new bone (5), hypertrophic cartilage (6), mature cartilage (7), or young cartilage (8) based on the characteristic heterogeneous mix of cell types, the dominant cell type, and extracellular matrix. As healing progresses the mix of cell types within a microscopic area changes. Some areas may undergo multiple tissue type transitions. A working hypothesis is that each of the microscopic tissue types is engaged in somewhat different activities, which are integral to the overall healing process.

The first stage 2 task was to select a square grid mesh size and overlay it on Fig 2A. Choice of mesh size was somewhat arbitrary. If it is too fine, there are fewer cells within the microscopic area and so the uncertainty in specifying the dominant cell type increases. If too coarse, the fraction of microscopic areas containing clearly distinguishable tissue types 4-8 decreases, rendering a single tissue assignment inadequate (and actions of the analog counterpart would likely require unique logic). A guideline for selecting grid size was that the cellular heterogeneity observed within the larger local callus area be reasonably preserved in the discretized image. For example, for a macroscopic region characterized by a heterogeneous mix of predominately of ~ 60% new marrow (gray) and ~ 40% osteoblasts (burgundy; new bone), the discretized image counterpart should be a mix of ~ 60% gray and ~ 40% burgundy tissue units. We selected a mesh size that corresponds to an 80×80 µm area in Fig 2A, which typically contained roughly 40 cells, and overlaid that grid on the day-7 and day-10i tissue sections. We then designated each microscopic area as being one of the nine tissue types. Fig 3 contains the resulting discretized, colorized images.

**Fig 3.**
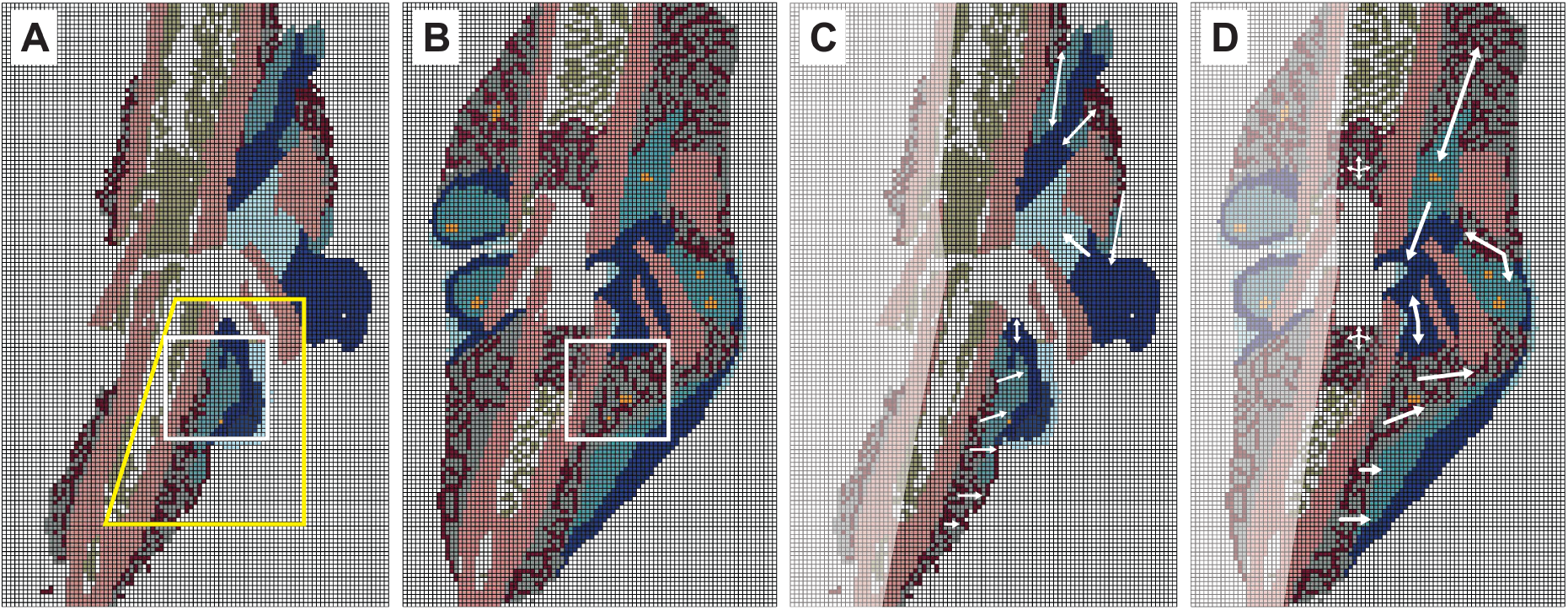
Discretized, images that map to sagittal sections through mouse tibia calluses. (A) Discretized, colorized counterpart to day-7 tissue section in Fig 2A. The yellow-box area corresponds to the one in Fig. 2A. The white box is the Target Region initial state. (B) Discretized, colorized counterpart of the day-10i tissue section. The white box is the Target Region final state. (C & D) Focus is drawn to the right sides. Arrows indicate directions of apparent local tissue changes. Some of that directional change that was ongoing in day-7 (C) continued into day-10 (D). Change in the other areas, particularly around the top edge of the Target Region, would have started after day-7. These apparent directional changes were taken into consideration for Mechanisms 2-4.

Although physically correct image interpolation (e.g., between day-7 and day-10i) is infeasible, sophisticated image interpolation methods, as demonstrated by Stich et al. [29], are available to create high-quality, convincing model images that represent unobserved transitions between recorded images of the same object. The criterion for an acceptable interpolation used by Stich et al., is qualitative: the interpolated images are perceived as visually correct by human observers. During stage 1, we faced the more daunting problem identified in Fig 4. We needed an image that plausibly anticipates the appearance of the mouse 1 fracture if it had been sectioned on day-10 rather than day-7. Starting with the features evident in Fig 2A, and drawing on the tissue features in Fig 2B, coauthor M.M. created an illustration of the envisioned mouse 1, day-10 section. It was judged plausible and acceptable by coauthor R.M. and, separately, by three independent domain experts (see Acknowledgements), thus concluding stage 1. Clearly, a different medical illustrator, one knowledgeable about callus progression, would create a somewhat different illustration. However, we suggest that variability introduced by such illustrations will not add measurably to the considerable variability and uncertainties already present, as illustrated by Fig 4.

**Fig 4.**
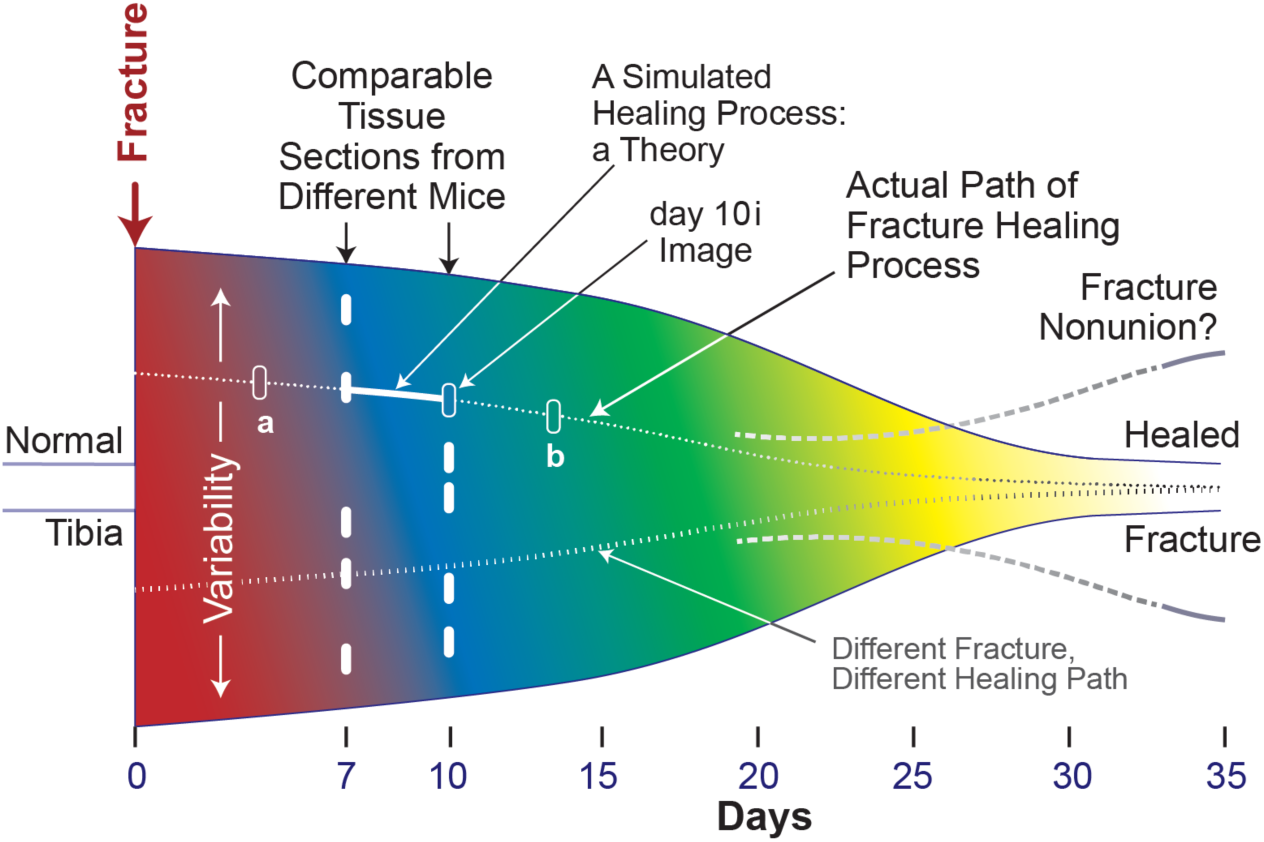
This is an idealized representation of key features of the space of tibia fracture healing in matched mice that are part of the same experiment. The height of the sketch represents the large interindividual variability, which is a product of each fracture being unique. Consequently, each healing process path is necessarily partially customized. Changing colors represent evolving subprocesses; the colors are not related to those used to represent the nine tissue types. The five short white bars on days 7 and 10 represent different tissue sections taken at comparable locations within different tibia fractures. The dotted path illustrates a trace of the healing process path for one of the five day-7 tibia fractures. Moving from day-7 to day-10 along the dotted path, the unshaded bar on day-10 indicates the illustration that we created, the day-10i tissue section. The two additional unshaded bars labeled a and b illustrate future extensions of the simulated Healing Process forward to day-14 and backward to day 4 *along the same healing path*. Two dashed gray lines illustrate that in humans fracture nonunion can occur. Absent interventions, nonunions in mice do not occur. The circumstances leading to nonunions in humans are unknown. The transition from one color to another occurs later toward the bottom, which illustrates that the pace of healing can be different from one fracture to another.

### Target Region

To demonstrate feasibility, we needed to select Target Region, but first, we needed to select a portion of Fig. 2A in which to locate the Target Region. For the latter, we selected the yellow-boxed area in Fig. 2A. It is bordered on one side by bone and marrow cavity, which means that transitions in that area will be focused rightward, rather than occurring in two or more directions. Because that area, and the corresponding region in Fig. 2B, exhibit similarities, we conjectured that the variety of feature changes occurring during transition from day-7 to day-10i might be representative of key healing features occurring elsewhere in the callus during that 4-day interval. Also, there is no indication that unique healing features may be occurring within this area but not elsewhere during that 4-day interval.

Limiting attention to just one target region can be viewed as a weakness. On the contrary, it is an essential part of a recognized, long-term mechanism-discovery strategy that can build on methodological lessons learned while using the Iterative Refinement Protocol in other contexts [22, 24, 26, 27, 30]. That strategy employs variations of the forward/backward chaining (described in S1 File) and requires selecting a Target Region (stage 3, Fig 1).

After we achieve stage 6 (Fig 1) for the day-10i Target Region (described below), we envision expanding the temporal reach of Callus Analog Mechanisms along the dotted line illustrated in Fig 4 to include achieving an earlier stage within that same Target Region, such as day-4i, and a later stage, such as day-14i, and doing so while continuing to simulate the original day-10i Target Region. Those objectives are illustrated by two unshaded bars labeled a and b in Fig 4. There are additional considerations. The histological evidence suggest that, on the same day, different subregions within a callus can be at different stages of repair and may be progressing at different rates. Given that, a parsimonious strategy is to select separated target regions within the same callus and develop simulations for each in sequence. They would be treated as independent modules. Future work based on simulations of independent target regions will help bring regional issues into focus prior to engineering their merger. The process of merging initially independent modules into a unified model of a tissue healing process would occur further downstream. Given that this work strives to establish the feasibility of the Fig 1 approach, it is efficient to focus first on just one Target Region.

Specifying the size of the target region is somewhat delicate. If the region is too large, with a large variety of tissue transition types, we run the risk that the process of discovering plausible and parsimonious logics to direct transitions will become unwieldy, possibly even problematic. If the region is too small, the variety of transition types may be too few to enable adequately simulating Target Region final state. We selected the 25×25 grid region designated by the white box in Fig 3A. Fig 3B shows the corresponding Target Region final state.

### Callus Analog Requirements

Simulation requirements—and thus software requirements—flow directly from desired use cases [31]. In the Introduction, we stated that a primary use case is exploratory simulations capable of the following: aiding image interpolation and providing plausible explanations for how callus features are progressively transformed, all while shrinking the space of possible explanatory transformation scenarios. The last two involve generation of plausible new mechanism-oriented explanations, as illustrated in Fig 1.

To realize use cases, we employ the virtual experiment approach described by Kirschner et al. [32] along with enhancements drawn from Smith et al. [27] and Petersen et al. [30]. In doing so, the methods employed must meet the following three requirements, which are based on broader sets of requirements discussed by Hunt et al. [31].

1. Achieve the two Fig 1 goals. To do so, requires that software mechanisms responsible for the transition of Target Region from initial to final state be plausibly biomimetic. We do that by employing the same type of software mechanism used successfully in other contexts [22, 25, 26, 27, 30] while adhering to the rigorous definition of mechanism provided in S1 File.
2. Ensure components, entities, and spaces (Fig 2) are concrete, acceptably biomimetic, and sufficiently modular during execution to facilitate analogical reasoning [33, 34].
3. Recognize that simulated healing arises mostly from quasi-autonomous component interactions at the lower level of Tissue organization.

To achieve Requirement 2, Callus Analogs are written in Java, utilizing the MASON multi-agent simulation toolkit [28]. The data presented herein along with Callus Analog code are available [35].

### Iterative Refinement Protocol

We customized the established Iterative Refinement (IR) Protocol [21, 26, 27, 30, 31] to meet the challenges evident in Fig 4. Given a software Mechanism that may explain a specified attribute and a virtual experiment design, the goal of an IR Protocol cycle is to test this hypothesis: upon execution, simulation features will mimic the target attribute within a prespecified tolerance. A concrete software mechanism can be falsified—shown to be inadequate—when, during the course of many Monte Carlo trials, it too often fails quantitatively to achieve its objective and/or exhibits non-biomimetic features. It is from encountering and overcoming such failures that explanatory insight improves. Each falsification improves credibility incrementally and shrinks plausible Mechanism space. Our customized IR Protocol follows:

1. State the objective for this IR Protocol cycle along with the target phenomenon to be explained. For this work, the latter is the day-10i Target Region.
2. Specify granularity of entities, activities, and spaces. Adhere to a strong parsimony guideline. Making the updated Mechanism too fine-grain can expand the space of plausible configurations beyond one's ability to manage efficiently.
3. Specify a similarity criterion. When first applying a quantitative similarity criterion (hereafter simply target similarity), make it weak initially, e.g., at least 40%, and increase it incrementally during subsequent IR Protocol cycles. Take small steps that often fail. For this work the final goal for simulated Target Region final state was that all simulated features be biomimetic (there are no non-biomimetic features) and exhibit ≥ 70% target Similarity. That value is sufficiently stringent, given the uncertainties illustrated in Fig 4. To illustrate, a slight change of grid placement can result in a 5-10% change in assignment of Tissue-unit (TU hereafter) types to grid-spaces. By doubling that range, we account for other sources of variability. With that in mind, a simulated final state that exhibits ≥ 70% Similarity to the day-10i Target Region supports the feasibility that the actual healing processes in that region and Analog processes are somewhat analogous at the current level of granularity.
4. When needed, revise Mechanism entities and activities. Record reasons for revisions. Revise in small steps. Frame the revision plan for this IR Protocol cycle as a hypothesis. An example: by making these changes to logic used by two Tissue Units, we will eliminate a non-biomimetic feature. Executing the revised Mechanism is thus a test of that hypothesis.
5. Conduct and evaluate many simulations—virtual experiments. Record and measure Target Region features each time step, noting when maximum Similarity is achieved. Each execution is a Monte Carlo trial of simulated healing. For this work, we studied multiple sets of 25 Monte Carlo trials.
6. **Failure**: one or more simulated healing features (e.g., patterns within the target region) are non-biomimetic and/or mean maximum Similarity < prespecified target Similarity. In this work, overcoming failure often required returning to step 4 and revising the Mechanism in some way. Failure provides new knowledge about model mechanism behaviors. Overcoming failure shrinks plausible Mechanism space. **Success**: go to step 7. With more mature model mechanisms, when appropriate, this would be the step at which knowledge of analog system behaviors would be expanded through similarity analyses and uncertainty quantification.
7. Choose one of these options. Return to step 3 and increase Similarity stringency. Return to step 2 and add a new similarity criterion. Add a new constraint, such as changing the logic used by a TU type when particular neighborhood feature is achieved and then return to step 4. Or, add a new phenomenon, such as this: gray and burgundy TUs form small clusters; and return to step 1.

### Callus Analog Mechanisms

Well-organized processes are responsible for the callus remodeling occurring between day-7 and day-10. Our operating hypothesis is that information available in day-7 and day-10 tissue section images can be used to draw simplified inferences about unobserved transitions occurring during intervening days, analogous to the approach used by Stich et al. in simulating unobserved transitions between recorded images of the same object [29].

A Callus Analog Mechanism as a system of biomimetic software entities and activities organized such that, during execution, it produces a phenomenon (a pattern) that is measurably similar to the day-10i Target Region. Feature changes within the Target Region explain how the pattern is generated. Stained tissue sections provide snapshots of the healing process. To be explanatory, a biological mechanism will exhibit the fourteen features identified in S1 File. Because Callus Analog Mechanisms exhibit those same features (also identified in S1 File), the two processes may be analogous.

Simulations are discrete time; time advances in steps. Fig 5 shows the Target Region initial and final state. The Process responsible for transitioning from initial to final state is the top-level Mechanism. Changes within local sub-regions from one time step to the next are lower level Phenomena. The lower level Mechanisms responsible for those changes are characterized by individual TU changes, which are controlled by the logic that governs TU actions during each time step. That logic is discussed below. During each time step, each TU, selected randomly, has one opportunity to update and act, based on changes that have occurred within its Moore neighborhood since it last updated. An action may change a TU type or one of its Moore neighbors. Coauthor R.M. identified the following as allowed but not required biomimetic transitions.

**Fig 5.**
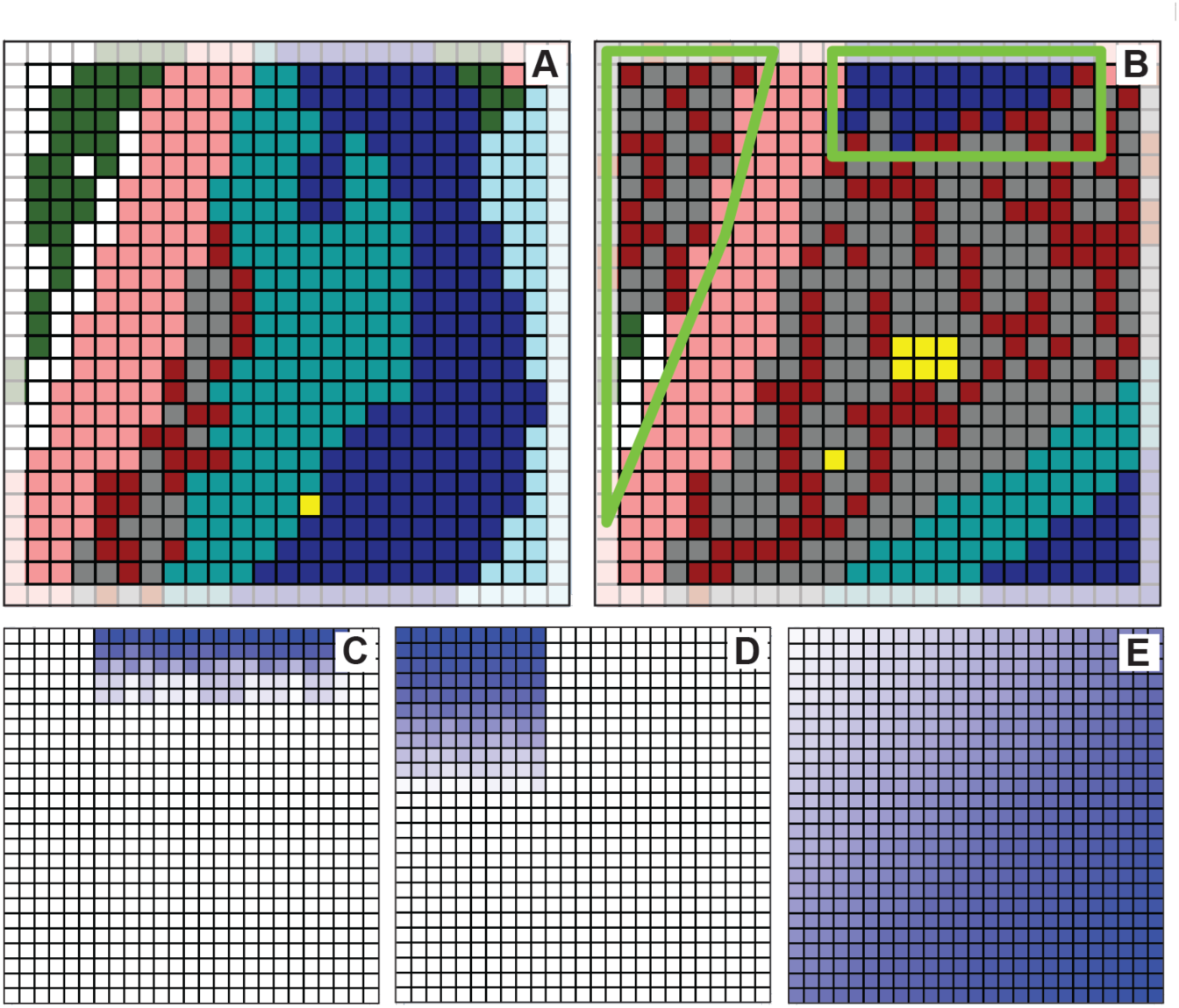
MASON displays of Target Region along with three influence grids. (A) Target Region initial state. (B) Target Region final state. The partially obscured TUs in the outermost columns and rows have an incomplete Moore neighborhood. They remained inactive for the duration of all simulations, but they do provide type information for their interior neighbors. All other TUs have 8 Moore neighbors and are active. For Mechanisms 2-4, behaviors of TUs within the two green-bordered regions require logic that is different from that used elsewhere. (C-E) The logic followed by each Mechanism 2 TU used probability values from one of these three grids. Darkest blue represents the largest value. White cells have a probability of zero or are unused. (C) Probability values used by TUs within the rectangular region. (D) Probability values hued by TUs within the triangular region. (E) Probability values used by TUs elsewhere.

- A marrow cavity (white, type 2) TU may transition into a new marrow (gray, type 4) or new bone (burgundy, type 5). New bone and new marrow form in parallel during callus maturation in a process that is indistinguishable. The formation of new bone and marrow occurs both outside and inside of the marrow cavity [9].
- Unidentifiable material (green, type 3) may transition into a mature cartilage (blue, type 7), hypertrophic cartilage (teal, type 6), new marrow (gray), or new bone (burgundy) TU. The marrow cavity often contains unidentified material in the earlier stages of healing. This material is replaced with new bone/marrow at later stages of the healing process.
- A hypertrophic cartilage (teal, type 6) TU may transition into a new marrow (gray) or new bone (osteoblasts, burgundy) TU. According to competing theories, hypertrophic cartilage is eventually replaced by new bone and new marrow in one of two ways, either by trans-differentiation [36, 37] or, as part of a two-stage process, they undergo apoptosis and are replaced by mesenchymal stem cells, which then transform into osteoblasts [35].
- A mature cartilage (blue, type 7) TU may transition into a hypertrophic cartilage (teal), new marrow (gray), or new bone (burgundy) TU. Hypertrophic cartilage develops from mature cartilage and eventually gets replaced with new bone/marrow. Because the time frame of the process in the mouse tibia is not clear, when comparing the day-7 to the day-10i Target Region, it may appear that mature cartilage has been replaced by either hypertrophic cartilage or new bone/marrow [39].
- A TU that is young cartilage (light blue, type 8) may transition into mature cartilage (blue). Cartilage cells in the callus undergo a maturation process from young cartilage cells to mature cartilage cells and then to hypertrophic cartilage cells [9, 39].

### Calculating Similarity

Each time step, the current simulated Target Region is compared to the Target Region final state and percent Similarity is calculated as follows.

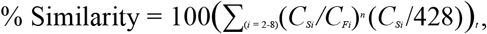

where *t* designates the time step, and *i* specifies the TU type, 2-8. *C*_*Si*_ is the count of TU type *i* in the <u>s</u>imulated Target Region; *C*_*Fi*_ is the count of TU type *i* in Target Region <u>f</u>inal state; 428 is the number of active TUs in Target Region; *n* = 1 if *C*_*Fi*_ > *C*_*Si*_, and *n* = –1 if *C*_*Si*_ > *C*_*Fi*_.

A case can be made that, if there is strong similarity between gray and burgundy TUs in the simulated and actual Target Region final states, then similarity scores should not be penalized because there are too many simulated gray TUs and too few simulated burgundy TUs, or visa versa. New marrow (gray) and new bone (burgundy) are always formed together. Thus, in some cases, the decision to designate an 80×80 µm area of stained tissue section as either gray or burgundy can be arbitrary; two experts may make different assignments. There are several ways to address that issue but they involve adding at least one new TU type. Given our strong parsimony guideline and the fact that we are at a very early stage in developing the Fig 1 approach, we elected to also calculate a Similarity value when gray are burgundy treated as the same during the calculation. The resulting value is designated Upper-Limit Similarity, simply UL-Similarity hereafter. A more realistic value may be between Similarity and UL-Similarity.

### Workflows

When working to discover plausible model mechanisms, there is a risk that the modeler, subconsciously or otherwise, will favor Mechanism features and logic that ensure that outcomes of generated behaviors are as the modeler thinks that they should be. We strove to eliminate that risk by adhering to the guidelines at steps 4 and 6 of the IR Protocol.

In developing the model mechanisms, we did not aim to include established biological features. Instead, we worked to develop model mechanisms that did not contradict known biology. Along the same lines, our model mechanisms were developed not to specifically describe characteristics of the fracture healing process, but instead to allow for a new, coarse-grained manner to think about the process. More information about advantages to avoiding absolute grounding can be found in [40]. By adhering to a strong parsimony guideline (IR Protocol step 2), we avoided adding unnecessary details; doing so enabled us to avoid inscription error. More details on overfitting and analog-to-referent mappings can be found in the Discussion section of Kim et al. [25]. In the subsections that follow, we describe four workflow stages, designated Mechanisms 1-4.

#### Mechanism 1

We sought logic that would enable each TU to transition from initial to final state. We limited attention to Von Neumann neighbors, the four TUs that share an interface. We identified 25 different types of initial-to-final state transition, along with six in which the TU at final state was the same as at initial state. Their frequencies are graphed in S1 Fig. For each transition type, we tabulated neighbors by type. We also tabulated how those neighbors transitioned from initial to final state. We used that information to devise conditions that must be met before a particular type of transition could occur. Each time step, with probability = 0.05 (chosen based on expected frequency of transition), each TU, selected randomly, was given an opportunity to act. When a random double from [0.0 - 1.0) was < 0.05, the TU stepped through its logic. Each TU transitioned to a final state only once. All transitions were completed within a few hundred time-steps.

We developed and tested a variety of rules. The following is an example of the two-stage logic structure, which was explored most extensively. Upon selection of a white TU, the random double is < 0.05, so it is given an opportunity to achieve its final state. It has three allowed transition options: white → gray, white → burgundy, or white → white. The above frequency data was used in advance to specify a probability for each option. Suppose that white → gray was selected. Specific conditions must be met in order for white → gray to occur. They are as follows. Are there ≤ 3 white neighbors, ≥ 1 green neighbor, or ≤ 2 gray neighbors? If yes, achieve final state by replacing self with a gray TU. If no, final state was not yet achieved. The vast majority of rules explored failed to enable the Target Region initial state to reach more than 50% Similarity to the Target Region final state. During time steps following that failure, one or more of its Von Neumann neighbors may transition. We explored allowing a TU that failed on its first attempt to have at least one additional opportunity to successfully transition during a subsequent time step, but that approach was abandoned when it became evident that many locations within the Target Region would need multiple transitions.

#### Mechanism 2

The healing process, which was well underway on day-7, created localized directional callus changes. White arrows on the right sides in Fig 3C and 4D indicate several apparent trends. Work on Mechanism 1 ignored those trends. For Mechanism 2, we hypothesized that the logic employed by all TUs should be location dependent. We used one of three probability grids to provide location dependent probability values. Several variants of the three gradients were explored. The three grids in Fig 5 were used in generating the largest Mechanism 2 Similarity values.

We posited that TUs within the rectangular region in Fig 5B behave differently from those elsewhere because they were being influenced by activities to their north, beyond the Target Region’s edge. They use the Fig 5C grid exclusively. Because TUs within the triangular region are isolated from other TUs by Bone, we posited that they would also operate independently; we reused the Mechanism 1 logic for that region but used the probability values in Fig 5D. Elsewhere, TUs utilized the Fig 5E grid. Probability values were employed as follows. Each TU normalized the probability values at the grid locations corresponding to its four Von Neumann neighbors. It then used the values to select one Moore neighbor for possible transition. Outside of the triangular and rectangular regions, gray, burgundy, teal, and blue TUs apply that logic each time step. TUs within that rectangular region operated similarly: a teal TU could be replaced by blue; a green could be replaced by gray or burgundy. As with Mechanism 1, each time step, with probability = 0.05, each TU was given an opportunity to act.

#### Mechanism 3

Mechanism 2 was insufficient. It exhibited non-biomimetic features. Rule details used in the final version were getting complex and thus deviating from our strong parsimony guideline. To enable achieving the targeted Similarity value and to produce a more biomimetic process, we inferred that each TU would need to be somewhat more fine-grained and use more location-dependent and goal-dependent logic. We retained the requirement that the probability to act each time step be the same (0.05) for all TUs. The Mechanism 2 directional probabilities were replaced by local vector values, which were refined during cycles through the IR Protocol. As an example, for the version of Mechanism 3 reported here, we specified that a blue TU could only cause a transition to the north, south, or east, and each direction was assigned a probability value. For blue TUs, those values were east = 0.6, north = 0.2, and south = 0.2. Once the location was selected, the TU type that would be the product on the transition was selected. The allowed transitions were those specified above. For example, a blue TU could transition to a teal or green TU but only when it had ≥ 4 blue Moore neighbors; otherwise, it remained blue. In Mechanism 3, green and white do not expand, but other TU types could generate them. Rules for all transition types are illustrated in Fig 7.

**Fig 6.**
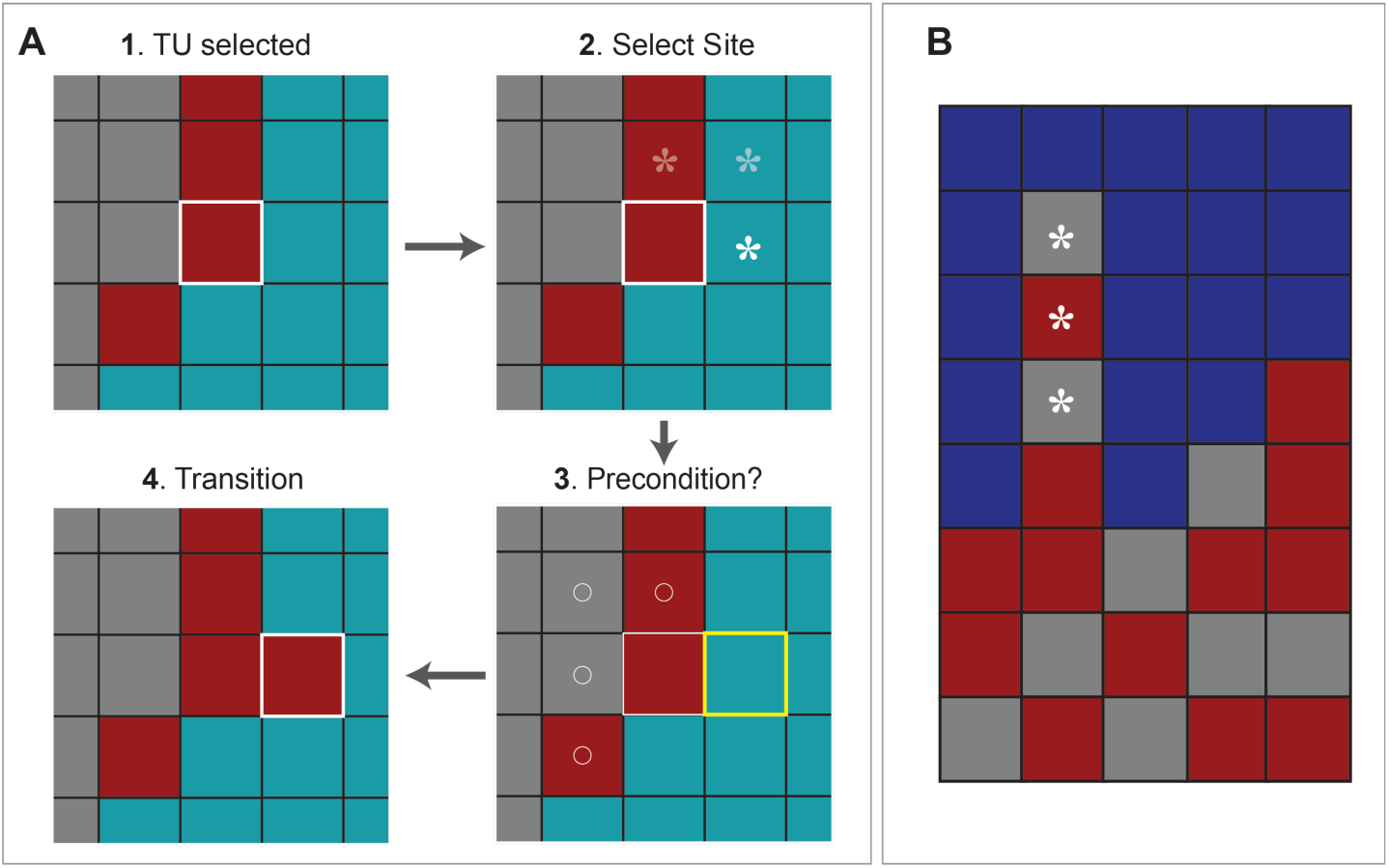
Selected Tissue Unit (TU) activities. (A) This sequence is an example of a sequence of events occurring within a single time step, and ending with a teal TU (outlined in yellow) transitioning into a burgundy TU. Step 1: the burgundy TU outlined in white is selected randomly. A probability determines if it is (or not) given an opportunity to act, which it is given in this example. It will not have another opportunity until the next time step. Thereafter, probabilities determine whether events at steps 2-4, in sequence, occur or not. The logic for all transitions is diagrammed in Fig 7. Step 2: a burgundy TU can initiate change in only one of the three TU neighbors marked by asterisks. With probabilities specified in Fig 7A, the East location is selected. Step 3: two questions are asked. Can the TU at that location transition? If yes, is the precondition for transition met? A teal TU can transition. Had the burgundy TU located north been chosen, the answer to the question would be no, because a burgundy is not allowed to transition. Nothing further would happen during that time step. The transition precondition for a teal TU is that the number of gray and burgundy TUs in the Moore neighborhood of the TU outlined in white be ≥ 4. In this example, the precondition is met (marked by circles). Step 4. A teal TU can transition to burgundy or gray with equal probability. In this example transition to burgundy occurs. (B) An example of a non-biomimetic feature (asterisks) occasionally encountered using Mechanisms 3 and 4 that requires a specialized rule, described in the text, to correct.

**Fig 7.**
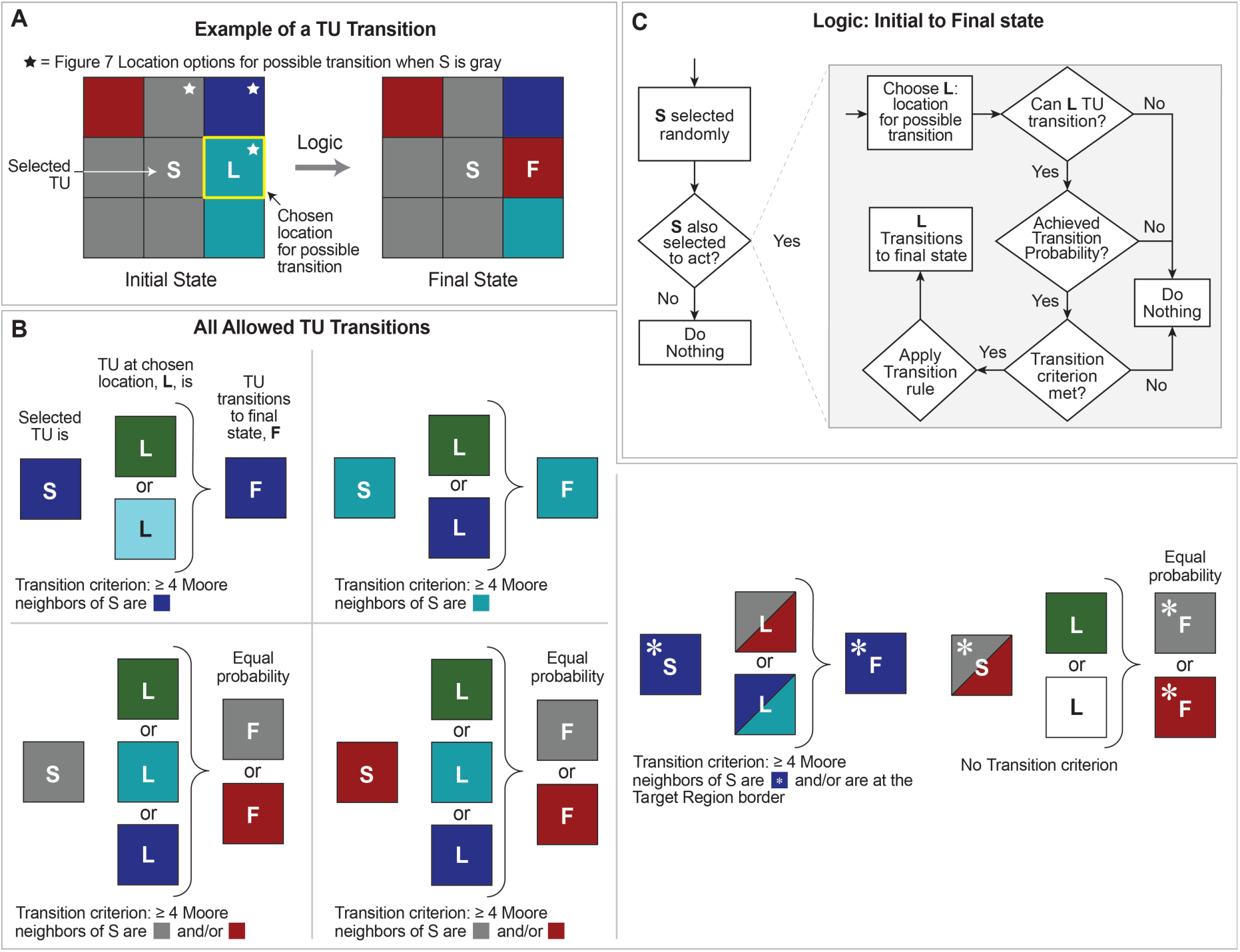
Tissue Unit (TU) Logic for Mechanisms 3 and 4. (A) This panel shows an example transition, which occurs within a single time step. A selected TU, S, identifies nearby locations for transition, selects a site, L, and, if the conditions further described in Figure 6A are met, L transitions to a new final state F. (B) Allowed transitions for TUs are shown. A selected TU, S, may initiate a transition for one of its neighbor cells, as determined by the probabilities shown in Fig 8. The probability grid determines whether a transition occurs for TU L, as well as if the conditions shown are met. For example, in the upper left panel, a blue TU is selected. It will next determine the site for transition, L, according to the probabilities in Fig 8. If L is green or light blue, L will be allowed to transition to blue. (C) Detailed logic for whether a transition occurs is shown. Details here are intended to expand upon Fig 6. At each time step, every TU is selected exactly once. Each TU is given an opportunity to act, according to a global probability. If selected, a TU, S, selects a site, L, for possible transition. Site L is selected based upon the probability grids in Fig 8 for the selected TU S. Whether L transitions is also governed by the probability in the center of the grid for S. If the probability is met and if L is one of the allowed types, shown in panel (B), L can transition.

It seemed clear from comparisons of Fig 3 sub-regions that, during the day-7 to day-10 interval, Target Region was being influenced by events to its north. We inferred that at sometime after day-7, expansion activity north of Target Region, illustrated by the white arrows in Fig 3D, would begin influencing the northernmost rows within the triangular and rectangular areas in Fig 5B. We explored several scenarios and selected this one for more thorough study: expanding TUs just north of Target Region would “collide” with Target Region TUs on each side of Bone, and a few of those invading TUs would actually enter the north edge of Target Region. After several cycles through the IR Protocol exploring that scenario, we observed more biomimetic simulated healing coupled with Similarity values ≥ 70% when the invading TUs used logic that was somewhat different from that used in Target Region, so we selected a particular version of that scenario and refined it further.

We needed a way to trigger the collision from the north. Given that the Target Region is a non-autonomous, smaller area of a greater whole. We considered specifying that it occur at some randomly selected time step during each Monte Carlo execution, but that risked increasing the variance for the time step with maximum Similarity. Our working hypothesis that a single, consistently applied TU logic could account for transition to the final state was falsified. Next, we tied the collision trigger to changes occurring within Target Region, which was intended to be reflective of changes occurring outside the Target Region. We specified that the collision be coincident with the first gray or burgundy TU reaching the northernmost row of active TUs, which we took as an indication that we had progressed sufficiently toward a simulated day-10 final state. Next, during that same time step, four blue invading TUs (blue*) were added to the central area of the rectangular region, three to the top active row and one to the row just below (variations on this theme work just as well). We use an asterisk to distinguish an original Target Region blue from an invading blue* TU. Although the TU types are the same—blue—blue* used somewhat different logic. During that same time step, a gray* and burgundy* TU are added left of center at the top of the triangular region.

A blue* TU is assigned its own logic. If it has ≥ 4 gray + burgundy Moore neighbors, it can expand. The direction is biased southward, as these cells have expanded from the north. The TU that is the product of expansion can transition into a gray, burgundy, teal, or blue TU. Gray* and burgundy* TUs also have their own logic. Their expansion is also biased southward. The TU that is the product of expansion transitions into a gray*, burgundy*, white, or green TU. All transitions are illustrated in Fig 7.

With introduction of blue* TUs, we encountered occasional unacceptable non-biomimetic features of the type illustrated in Fig 6B. We inferred from Fig 3 that blue TU sub-regions should expand and contract somewhat cohesively. However, as blue TUs expand downward, they may encounter an obstacle—a column or “wedge” containing gray, burgundy, and/or teal TUs. We observed such obstacles persisting. That persistence “splits” the blue sub-region, breaking the apparent cohesion. To prevent such occurrences, we added an additional rule: a gray, burgundy, or teal TU with ≥ 5 blue Moore neighbors replaces itself with a blue TU. After including that rule, the average maximum Similarity value for Mechanisms 3 improved.

#### Mechanism 4

Values governing Mechanism 3 location selection probabilities were hard-coded for each TU type. Mechanism 4 was the result of work to address Requirement 2 by increasing flexibility and streamlining Mechanism 3 activities. Each of the seven active TU types was assigned a mini-grid of expansion probabilities, illustrated in Fig 8, which it applied to its Moore neighborhood. Mini-grids are easily customizable, and are provided to the Analog through standard comma-separated values (CSV) files. The Mechanism 3 rule that prevented splitting of an expanding blue sub-region was carried forward without change. Simulated Target Region final states generated during executions of Mechanisms 3 and 4 are statistically indistinguishable.

**Fig 8.**
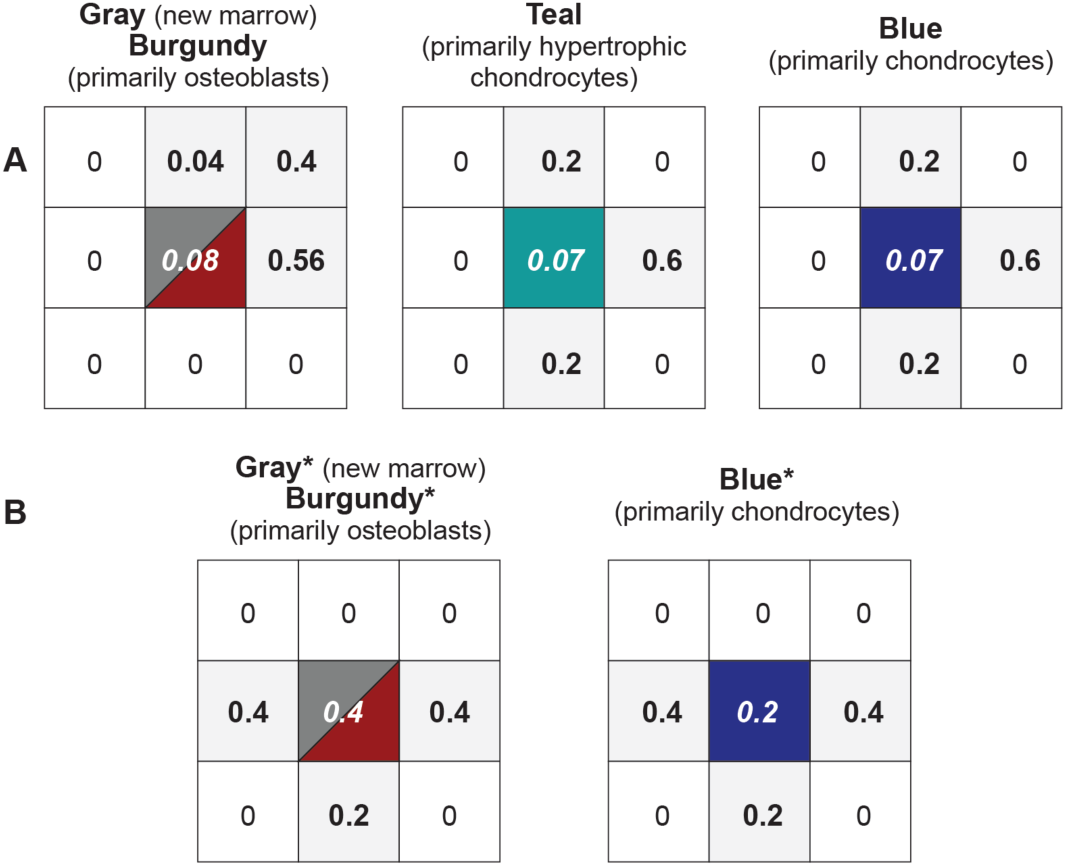
Probabilities governing TU transitions in Mechanisms 3 and 4. The center value is the probability each TU will have an opportunity to act each time step. The value at each shaded location specifies the probability that a transition event may occur at that location. The types of allowed transitions are illustrated in Fig 7. (A) The precondition for transition is that each of these four TU types have ≥ 4 Moore neighbors that are of the same type. (B) These are the three TU types that enter Target Region from the north. Their rules are different from counterparts in A. Gray* and burgundy* have no precondition for transition. They can transition into gray*, burgundy*, white, or green. The precondition for a blue* transition is that ≥ 4 Moore neighbors are blue* and/or are at the Target Region border. It can transition into gray, burgundy, teal, and blue.

### Internal Control

What further improvements in similarity values, as calculated above, might reasonably be achieved? We answered that question with Mechanism 4 internal control calculations that draw on the fact that the closest Similarities that can be achieved for a simulated day-10 Target Region will be those achieved by independent executions of the Mechanism that generated it. The analog Healing Process from day-7 initial state to a simulated day-10 final state is unique for each Monte Carlo execution of Mechanism 4. We selected one Mechanism 4 Monte Carlo execution from 25 and recorded its Target Region configuration at the time step for which simulated Target Region final state maximum Similarity was achieved. We designated it to be the internal control simulated day-10 target state. We then measured maximum Similarity of each of the other 24 Monte Carlo Healing Processes to that simulated day-10 target state.

### Data Types, Reuse, Code Availability, and Sharing

Callus Analogs are a form of data, using both the implicit schema of MASON/Java and the explicit schema of configurations. Analog and configuration data are maintained, archived, and released using the Subversion version control tool in two repositories: one public for collaboration and one private (Assembla) for rapid and prototyping development with project partners [35, 41]. Input-output (I/O) data is handled separately. Smaller data sets (tissue data) are stored in simple CSV format.

The entire Callus Analog toolchain is open-source, thereby enabling repeatability. Similarly, all generated and released data from the project is licensed and available as open data. Callus Analogs are built for, maintained, and, when needed, can execute in a cloud environment (e.g., Google Compute Engine) to ensure platform and infrastructure repeatability across future experiments, project team members, partners, and the wider community.

### Adherence to Simulation Best Practices

The protocols, procedures, and methods that we employ to insure that results of Callus Analog experiments are reproducible and to build credibility that they are scientifically useful are described in detail in [27]. Except as noted, we followed those best practices during the workflows described above. They include 1) quality assurance and control protocols, 2) establishing face validation, 3) verification procedures for model mechanisms, 4) ensuring repeatability, 5) methods for generating narrowly focused predictions, and 6) Callus Analog validation methods used during IR Protocol cycles.

However, it is too early for systematic sensitivity analyses or efforts to quantify uncertainties associated with particular Mechanism 4 configurations. That is because Callus Analog is still at a very early stage; the focus is on acquiring new insights. Mechanism life cycles are expected to remain relatively short. As soon as we target additional attributes, it is likely that Mechanism 4 will be falsified (because it cannot achieve those new targets). Thus, it will be necessary to alter Mechanism 4 during new cycles through the IR Protocol. We acquire evidence that we are adhering to our strong parsimony guideline at IR Protocol step 2 in part through documentation of sources of uncertainty and focused assessments of simulated final state sensitivities to modest changes in the logic followed by each TU. An example is provided in Results.

## Results

Each execution of Mechanisms 2-4 is a unique, simulated healing process, which may (or not) mimic an interval of actual fracture healing. Because the focus is on similarity to the day-10i Target Region, we refer to the simulated state having maximum Similarity value for a given Monte Carlo execution as the simulated day-10 Target Region final state. However, we currently have no data to guide mapping time steps to wet-lab clock time.

Summarized maximum Similarity results for Mechanisms 2-4 are included in Table 1 and the complete results are included in S1 Dataset. Examples of simulated Target Region final states for Mechanisms 2-4 are provided in Fig 9. S1 Video is an example of the complete simulated healing process for Mechanisms 2. It includes the final state having the largest Similarity value. Mechanism 2 improved upon Mechanism 1, generated reasonable simulated states, but failed to meet the biomimesis requirement. A feature of a simulated healing process that has no known real counterpart is designated non-biomimetic. Fig 9A shows the simulated Target Region having the largest maximum Similarity value from a set of 25 executions. Although maximum Similarity values of over 70% were achieved for Mechanism 2, at least two non-biomimetic features were observed. 1) In simulated final states, teal TUs were absent from the lower right region of Target Region and that reduced Similarity values significantly. The small islands of teal and blue TUs within a gray/burgundy region in Fig 9A may also be non-biomimetic features. 2) Temporal profiles of Similarity values (not shown) reached a plateau prior to time step 5000, which persisted beyond time step 20,000. Consequently, the time step at which maximum similarity occurred appeared somewhat random.

**Table 1.**
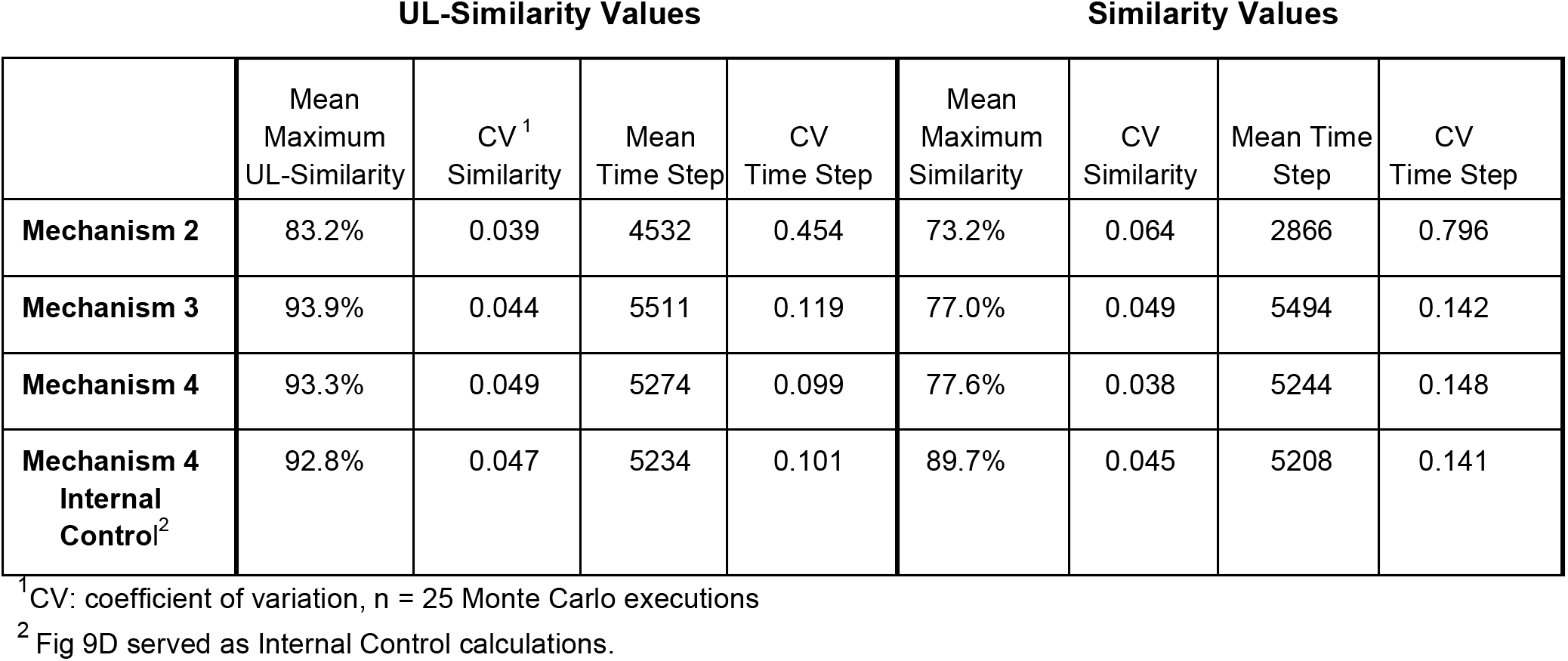
Similarity Criteria

**Fig 9.**
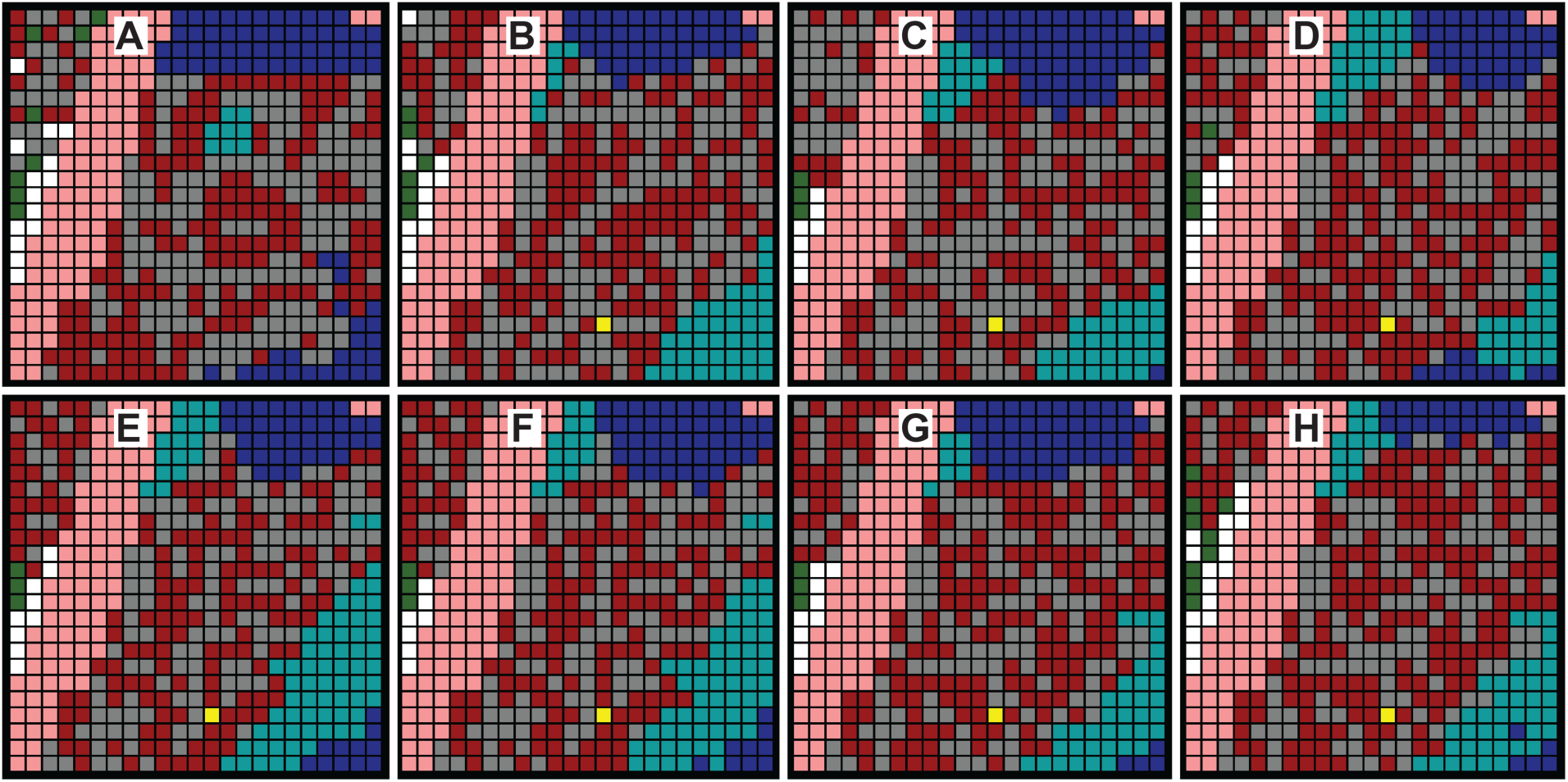
Examples of day-10 simulation results. From the sets of 25 maximum Similarity values (one for each Monte Carlo execution) from which the Table 1 values were derived, we selected these eight examples. The first four have the largest maximum Similarity value for the specified set: (A) Mechanism 2, (B) Mechanism 3, (C) Mechanism 4 when calculating Similarity, and (D) Mechanism 4 when calculating UL-Similarity. (E-H) From all other Mechanism 4 maximum Similarity values, we selected these four as exhibiting good overall biomimicry to the day-10i image based simply on visual comparisons. E and F are from the Table 1 set when calculating UL-Similarity. G and H are from the set.

Fig 9B shows the simulated Target Region having the largest Mechanisms 3 maximum Similarity value from a set of 25 executions. S2 Video includes Fig 9B. Table 1 data shows that Mechanism 3 improved on Mechanism 2. However, the absence of blue TUs adjacent to teal in the lower right limited maximum Similarity values and may be a non-biomimetic feature.

Figs 9C and 9D, which are included in S3 and S4 Videos, are examples of simulated Target Regions having the largest Mechanisms 4 maximum Similarity value from the set of 25 executions summarized in Table 1. No non-biomimetic features were observed. Based on simple qualitative visual comparisons, we judged the similarity of Figs 9E-H to the day-10i Target Region to be comparable to that of Fig 9C,D, even though their Target Regions had smaller maximum Similarity values. That observation indicates that, moving forward, improved measurements of Similarity will be needed. S5 and S6 Videos includes Figs 9E,F.

Each Mechanism 4 is a unique, simulated healing process, which is intended to mimic the interindividual variability of actual fracture healing. To observe and measure that uniqueness, single Mechanism 4 executions were selected from those used to provide the summary results for Mechanism 4 in Table 1. A Similarity value was calculated and plotted each time step in Fig 10.

**Fig 10.**
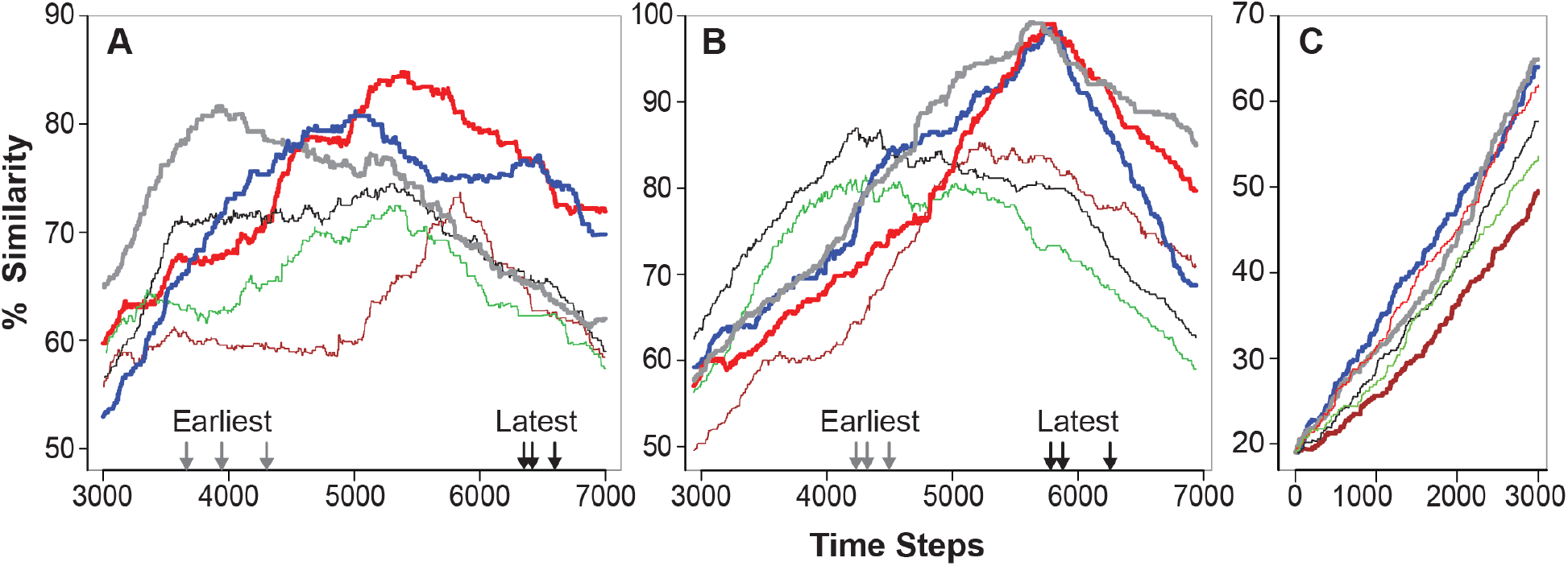
Temporal profiles of Similarity values for selected executions. Single executions were selected from those used to provide the summary results for Mechanism 4 in Table 1. (A) Profiles selected from the 25 Monte Carlo executions when calculating Similarity. The three profiles with the wider line widths exhibited the largest maximum Similarity values. The three profiles with narrowest line widths exhibited the three smallest maximum Similarity values. Arrows at bottom: those on the left mark the three earliest occurrences of a maximum Similarity value. Those on the right mark the three latest occurrences of a maximum Similarity value. (B) Profiles selected the 25 Monte Carlo executions when calculating UL-Similarity. The presentation is as described in A but y-axis values are different. (C) Examples are plotted of ascending portions of Similarity value profiles when calculating Similarity. Two of wider profiles exhibit the steepest increase. The wide profile at bottom exhibit the slowest increase. The three narrow profiles were selected randomly from the other 22.

Looking for such features and for wet-lab evidence that pushes the decision either way should be part of future research Callus Analog research. We saved the images corresponding to the maximum Similarity for both sets of 25 Mechanism 4 executions summarized in Table 1. Similarity value is just one way to judge good overall biomimicry.

Assessments of simulated final state sensitivities to modest changes in TU logic help identify sources of uncertainty. They also provide evidence for how tightly we are adhering to our strong parsimony guideline. For example, changing action and event probabilities in Mechanism 4 by ± 10-15% for a particular TU produces changes in simulated final state and temporal profiles of Similarity values that are well within the range produced by 25 Monte Carlo executions; the behavior space of Mechanism 4 is not significantly altered. The following is a specific illustration. We changed the Moore neighborhood probability values for blue (randomly chosen) in Fig 8A from [0.2 north/south and 0.6 east] to [0.165 north/south and 0.67 east] and repeated the 25 executions tabulated Table 1 using the same seeds. The average maximum UL-Similarity was 93.6% vs. 93.3% in Table 1; and for Similarity it was 77.8% vs. 77.6% in Table 1. For mean time step at which those value occurred, the new (vs. Table 1) time step was 5317 (vs. 5274) for UL-Similarity and 5299 (vs. 5244) for Similarity. However, changing how and when invasion of TUs from the north is triggered is an example of a change that can have a more significant influence: for such changes, the behavior space of Mechanism 4 can be significantly altered.

## Discussion

Given the reality illustrated in Fig 4, the prospect of discovering plausible mechanism-based explanations of fracture healing by relying solely on results of wet-lab experiments is problematic. We sought computational methods that could be facilitative. We ruled out established biomedical multiscale modeling and simulation methods, as recently reviewed by Walpole et al. [42]. A requisite for the latter is having sufficient information and knowledge to provide an adequately detailed mechanism-based explanation that is believed to account for the phenomenon of interest. That requisite can be met when the focus of the research can be characterized as being right-of-center on the four Fig 11 spectra. Each spectrum represents a broad attribute of the research. Location on the spectrum characterizes current reality. Fracture-healing research cannot yet meet that requisite because it is characterized by locations on all four spectra that are considerably center-left. For center-left locations, combining analogical models (e.g., electrical, mechanical, systems engineering, etc.) and analogical reasoning is a proven alternative strategy for developing plausible explanations of phenomena [33, 34]. But because there are no material systems that can serve as analogical models, we are focusing on developing model mechanisms in software.

**Fig 11.**
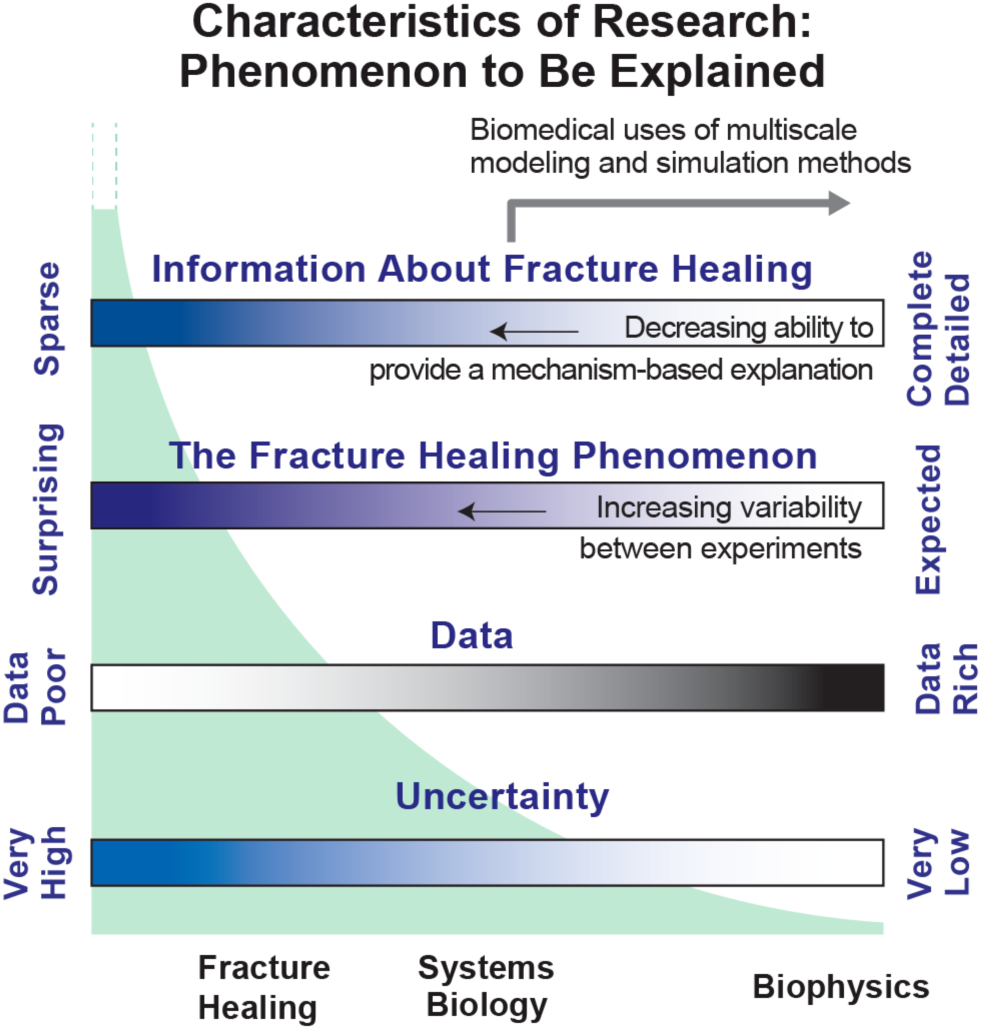
Characteristics of obstacles listed in the Introduction. Each spectrum provides a different perspective on research aimed at characterizing and explaining a phenomenon, fracture healing in this case. Relative to research in other biomedical domains, fracture-healing research can be characterized by a left-of-center location on each spectrum. Spectra locations considerably right of center are most supportive of the conventional inductive methods used by current multiscale modeling and simulation researchers [45]. The green shaded curve illustrates that, as one moves leftward, the number and variety of equally possible explanations increases dramatically, which can be a serious barrier to progress.

The issues that are the focus of fracture-healing research are located center-left on the four spectra. Consequently, there are many equally possible explanations for the healing phenomenon. That reality is illustrated in Fig 4 and by the green curve in Fig 11. The conventional strategy is to perform new experiments that generate new data and new knowledge. In doing so, one’s location shifts rightward and that shrinks the number and variety of equally possible and plausible explanations. Given the Fig 4 reality, that approach is not available. As explained in [21] and demonstrated in [22, 23, 25, 27], an advantage of the software-based model mechanism approach is that, by keeping model mechanism entities parsimoniously coarse grain and concrete, we limit the variety and space of model mechanisms capable of generating the targeted phenomenon.

By showing that Mechanism 2 was inadequate, we eliminated it from further consideration along with all finer grain variants of Mechanism 2. By so doing, we shrank the space of possible model mechanisms. In moving from Mechanism 2 to 3, the granularity of the concrete model mechanism entities is unchanged, but their activities are changed. Thus, we are working within a marginally smaller possible model mechanism space. As demonstrated in [27], we can reduce the space of possibly explanatory model mechanisms more significantly by increasing the number and variety of targeted phenomena—validation targets—that require the analog system to satisfactorily simulate, and that is our plan for moving forward.

The effort to keep the Callus Analog simple is where we encountered conflicting demands. To be scientifically useful and facilitate discovery, the Callus Analog must be sufficiently biomimetic in both model mechanism and generated phenomena. Increasing biomimicry requires that we make Callus Analog features finer-grain. However, making a Callus Analog finer-grain absent an evidence-based reason for doing so (as stipulated by the IR Protocol) risks dramatically increasing the space of equally possible explanatory mechanisms. Adhering to a strong parsimony guideline at step 2 of the IR Protocol helps resist that pressure.

### Simulated healing

Each Callus Analog execution provides a record of an analog Healing Process, which is the top-level analog phenomenon. Executions generate the succession of changes by which earlier states of the Target Region gradually become a simulated Target Region final state. Each video explains how the initial state is transformed into a final state that is measurably similar to Target Region final state. It is too early to claim that strong analogies actually exist between temporal features of simulated and actual fracture healing for Mechanism 4. Nevertheless, we can draw new inferences, which can be investigated, and/or directly tested (in some cases, supporting/refuting evidence may already exist but has not been connected to the new inference). We can hypothesize that, at comparable levels of granularity, the simulated healing processes seen in the S3-S6 Videos have actual fracture healing counterparts. Suppose that we accept that hypothesis: the Mechanism 4 healing process and the actual corresponding fracture healing are strongly analogous. Given the logic detailed in Figs 7 and 8, what might one infer about the intra- and intercellular activities ongoing within (and between) each 80×80 µm area of callus tissue (corresponding to a Callus Analog TU)? Even though the cellular components within each area are typically heterogeneous, we would infer that their activities are tightly orchestrated. Further, continuously updated information about activities within neighboring tissue areas in all directions is needed to maintain those orchestrated, goal-directed activities. Simultaneously, those same cells are providing information about their own activities to neighboring tissues. When current information exhibits specific features, the cellular components signal the dominant cellular component within a particular neighboring tissue area to change its behavior in a particular way, i.e., initiate a transformation (e.g., hypertrophic chondrocytes in particular area are signaled to transform into osteoblasts). The different neighboring regions providing influential information may span a cross-sectional area as large as 240×240 µm (maps to a TU’s Moore neighborhood). Such information processing and distribution would require sophisticated, robust intercellular communication networks. However, there is no current evidence that such networks exist. On the other hand, there has been no reason to date to look for such evidence.

Taken together, the S3-S6 Videos are low-resolution (coarse-grain) models of explanation— theories—that maps to a 4-day portion—day-7 to day-10—of the tibia fracture healing process in a mouse. There are currently no comparably detailed competing theories of fracture healing. Because Callus Analog mechanisms are concrete, they are easy targets for scientific challenge, and it is through that use that we anticipate Callus Analogs will provide scientific value moving forward.

The changes occurring within the Target Region during Mechanism 4 executions are intermediate level phenomena; they have histomorphometric counterparts in callus tissue sections. Two examples are the eastward progression of teal TUs replacing blue TUs and the eastward expansion of the mixture of gray and burgundy TUs. The mechanisms responsible for those intermediate level phenomena are mediated by the individual activities of the participating TUs. The logic dictating TU actions at each time step, as illustrated in Fig 7C, provides Mechanism orchestration. A change in TU type at a particular grid location is the lowest level (finest-grain) model mechanism phenomenon.

### Iterative Mechanism improvement

Mechanisms 1-4 were developed sequentially. We derived the most useful results when a particular configuration (logic, utilization of neighborhood information, probability values, etc.) failed to meet expectations. Some failures were marked by poor maximum similarity values. Others were marked by a feature within Target Region that was unexpected or judged non-biomimetic. In such cases, we would hypothesize a plausible explanation for the problem and a possible solution, and then conduct experiments to challenge that hypothesis and solution. When successful, we achieved an incremental Analog improvement. Failures provided new knowledge by enabling us to marginally shrink the space of plausible biomimetic healing processes. Mechanism 1 was unsuccessful because it was flawed in several ways. Nevertheless, observations made during IR Protocol cycles stimulated ideas for other logic that might be explored, including the ideas that drove development of Mechanism 2 and, later, Mechanism 3.

From observations made during explorations leading to Mechanisms 1 and 2, we inferred that, in order to make simulated healing processes more biomimetic, it would be necessary to include two features. 1) Allow for multiple TU changes at any grid location during simulated healing. 2) Have sustained directional influences, spanning many TUs, guiding or constraining the direction and type of TU transitions. The latter may map to the combined net effects of multiple factors such as relative abundance and activity of immune cells [43] signaling influencing osteogenic and chondrogenic transcription networks [44], O_2_ gradients [45, 46], and mechanical influences [47]. Callus Analog has achieved its current objectives without needing to bring any of those influences into focus, consistent with our strong parsimony guideline. As the list of targeted phenomena expands, it will become necessary to make the model mechanism finer-grain. It is during such refinements that a newly added Callus Analog feature may map to one or more of those influences.

The logic used by Mechanisms 3 and 4 limits the direction in which a TU can affect transition of a neighbor, and it imposes preconditions on number and type of Moore neighbors that must be present before a TU transition can occur. A consequence of those goal-directed constraints was emergence of apparent cohesion of TUs within three areas that are clearly evident in Fig 9E-9H: one area dominated by blue TUs, another dominated by teal TUs, and a third dominated by gray and burgundy TUs. That apparent cohesion is clearly evident during S2-S6 Videos. From the simulation engineering perspective, Callus Analog could be simplified if those three areas were represented as large, quasi-autonomous, sub-Callus organized units, within which TUs are simply parts under the control of each organized unit. However, there is, as yet, no direct wet-lab evidence that would support or require such simplification.

Interfaces between those areas map to well-documented transition zones (e.g., see [39]). Moving forward, features of transition zones will be added to an expanding list of targeted phenomena to further shrink the space of plausible explanatory model mechanisms.

Special attention was given to understand why, during an IR Protocol cycle, a model mechanism failed. As Petersen states, "having that information is essential to the scientific process because it is falsification that provides new knowledge: specifically, the current (falsified) mechanisms are flawed—they are not a good analogy of the referent biological mechanisms" [30]. Building upon and revising flawed hypotheses offers a new perspective and new way of thinking about plausible networked callus healing processes, and that new way of thinking may well become the primary value of the Callus Analog approach.

### Current competing theories

Fracture healing occurs primarily through the process of endochondral ossification, a process in which cartilage matrix is replaced by bone. This is the same process by which many bones are formed and grow. During endochondral ossification at the fracture site, chondrocytes express vascular endothelial growth factor, which induces vascular invasion of the cartilage matrix [48–50]. Along with the invading vasculature, osteoclasts arrive at the callus and degrade the cartilage matrix. Previously, the chondrocytes were thought to undergo programmed cell death [38], and concurrently, osteoblasts, which are delivered by the vasculature [18], replace cartilage matrix with bone. In this two-stage theory, chondrogenesis—cartilage development—serves chiefly as a means for producing hypertrophic chondrocytes, and they, in turn, initiate bone formation (carried out by other cells). However, a competing theory has emerged. Chondrocytes enter a transient stem-cell-like state from which they transform into the osteoblasts and osteocytes that form the new bone [36, 37]. The earlier theory envisioned chondrogenesis and osteogenesis as separate processes, whereas the more recent theory is mechanistically simpler; it envisions chondrogenesis and osteogenesis as characteristic sequential features of the same process.

All of the wet-lab observations on which those two competing theories are based are below the resolution of Mechanism 4. They are subsumed by the Fig 7 logic. So, at this stage, we have no evidence for or against either theory. As we add new target attributes downstream, we envision replacing the Fig 7 logic with model mechanism details using the tuneable resolution process of Kirschner et al. [32] (described below), and at that stage we should be able to challenge those competing theories.

### Strengthening weaknesses; addressing limitations

Because we are still early stage, the Mechanism 4-based simulated healing process comes with an ample supply of weaknesses and limitations. Both a weakness and limitation is that there are no (fine grain) 1:1 counterparts to the cellular entities and molecular level events that are the focus of the majority of wet-lab experiments. As Callus Analog credibility improves and finer-grain features are included as validation targets, it will become feasible to increase model mechanism resolution further utilizing Kirschner’s tuneable resolution process [32], which is a systems biology approach for discretized multi-level, multi-compartment computational models. The process involves fine- or coarse-graining of entities and activities. Such an approach allows for the adjustment of the level of resolution specific to a question, an experiment, or a level of interest.

The day-10i illustration is a stage 1 requirement. It is an important source of information but also a source of uncertainty. A requisite for building an explanation for fracture healing is having staged representations of the same fracture (e.g., on days 4, 7, 10, 14, 20, etc.) that are, within reasonable tolerances, reliable, semi-quantitative, and scientific. There are currently no protocols to achieve that requisite. However, many histologists, pathologists, and biologists are trained in accurately illustrating representations of specimens, including tissue sections. A logical next step would be to acquire two independently generated day-10i illustrations of the same section and then document where and why they are similar and different. Thereafter, we envision protocols and methods for developing credible staged illustration representations becoming increasingly standardized, and where feasible, automated. To increase standardization, we can draw from the robust best practices developed over decades for standardization of pathologic and histologic evaluations and reporting. We can also draw on the medical image registration methods [51] that enabled rapid advances in computed tomography and magnetic resonance imaging.

Feature discretization and simplification at stage 2 helps manage uncertainties, but the process itself is also a source of new types of uncertainty. There is a risk that increasing or decreasing grid mesh density can alter analog-to-mouse healing process mappings in scientifically meaningful ways. A good future time to assess that risk will be when Callus Analog insights have advanced sufficiently to begin exploring for the first testable theory for fracture nonunion.

The cellular components of all 80×80 µm areas within the day-7 tissue section Target Region are heterogeneous, but discretization requires that the corresponding analog grid space be occupied by one of nine TU types. To limit unintended bias, we can, as above, continue to draw from the mature best practices of pathologists to develop protocols to minimize added uncertainties. Longer term, we envision discretization protocols becoming automated. Near-term, several strategies can be explored to discover and ameliorate discretization weaknesses. Here are two examples. 1) Acquire two (or more) independently generated target region discretizations, and then independently develop an analog for each that achieves the same final state Similarity criteria. 2) There will always be instances where it will be difficult for a domain expert or an automated process to make some TU assignment, such as choosing between a new marrow TU (gray) or an osteoblast-dominated TU (burgundy), because those cell types are similar. In such cases, those grid spaces can be given a new designation, TU = g/b. At the start of each simulation experiment, all g/b spaces are randomly designated as either gray or burgundy. The result is a set of Monte Carlo Target Region starting states. One then cycles through the IR Protocol for each in parallel until the Target Region final state Similarity criterion is achieved. Both strategies require increased work, and automating IR Protocol tasks will help avoid reducing the overall workflow pace.

Selecting an initial target region at Stage 3 was essential to demonstrate feasibility. Moving forward, the approach must be expanded in stages to cover the entire callus, possibly as follows. First, develop and improve simulated healing processes for small portions of a callus and then explore how best to merge them incrementally to simulate more of the fracture-healing phenomenon. It seems likely that multiple sub-callus processes will be needed to simulate healing within the entire callus. The evidence suggests that different callus subregions can be at somewhat different stages of repair and may be progressing at different paces. Separate simulations of independent sub-regions will help bring these issues into focus. A plausible next step would be to select a new day-7 target region (possibly larger than 25×25) and determine if Mechanism 4 is able to achieve a corresponding day-10i final state with Similarity values > 70%. Following that, we envision extending those two analog healing processes forward to day-14 and backward to day-4.

Each Mechanism 4 video is a sample from the circumscribed space of Mechanism 4-based model healing processes. Each is biomimetic. Are all other Mechanisms in that space also biomimetic? It is too early to answer, but it seems likely that the answer is no. Each video (the record of one Monte Carlo trial) provides a means to search for and address the emergence of non-biomimetic features. Observing more videos provides one with a better overall impression of the space of simulated healing. Domain experts observing videos can identify features that may be non-biomimetic, such as the small islands of teal and blue TUs within a gray/burgundy region in Fig 9A cited above. Features that appear in one video may not appear in another. Assume that domain experts identify a likely non-biomimetic feature in several Mechanism 4 videos. Mechanism 4 would be falsified. We would then seek a marginally different—yet still parsimonious—model mechanism in which the logic has been revised to avoid exhibiting the non-biomimetic feature and still meets all similarity criteria. The revised Mechanism would circumscribe a *smaller set* of analog healing processes. In the preceding scenario, the videos provide domain experts a new means of thinking about callus healing. More broadly, simulated healing provides a new perspective on the actual healing process, and it is from that perspective that we encourage the use of simulations to enhance the mechanism discovery process.

## Acknowledgements

This work supported in part by NIH-R01AG046282 (R.M.) and the UCSF BioSystems group (C.A.H). We thank domain experts Louis Gerstenfeld, Dana Graves, and Kurt Hankenson for validating the day-10i illustration. We also thank Glen E.P. Ropella and Andrew K. Smith for constructive criticism.

## Author contributions

R.C.K., M.M. and C.A.H. conceived and designed the experiments; R.C.K. performed the experiments; R.C.K. and C.A.H. analyzed the data; R.C.K., C.A.H. and M.M. wrote the paper; R.C.K. and C.A.H developed and verified Callus Subregion Analogs and sub-component requirements; R.M. and M.M. guided refinements of analog to referent mappings; M.M. contributed to targeted attribute selection and Similarity Criteria development; R.C.K. complete IR Protocol cycles; R.C.K. developed scripts for data analyses and data visualizations; R.C.K., C.A.H., M.M. and R.M. contributed manuscript content; R.C.K., M.M., and C.A.H. guided the research.

## Supporting Information

**S1 Fig. Frequency of Allowed Transitions used in Mechanism 1 Development.** This plot depicts the frequency of the 25 allowed transition types, which include transitions where the final state was the same as the initial state.

**S1 File. Discovering Biologically Explanatory Mechanisms.** Forward/backward chaining reasoning strategies are described. They aid discovery of explanatory model mechanisms. Moving forward, we will rely on those strategies to expand Callus Analog to earlier (e.g., day-4) and later (e.g., day-14) stages of fracture healing. Mechanism is defined. To meet the definition of mechanism, the software mechanisms must exhibit the features listed in S1 Table. The table describes features that should be exhibited by an explanatory mechanism, along with the Callus Analog counterparts.

**S1 Dataset. Raw Data for Results.** All data reported in the manuscript are provided in the spreadsheet. The spreadsheet is organized by tabs. The “Statistics” tab contains the summarized data provided in Table 1 of the manuscript. Similarity and UL-Similarity data are provided for the Monte Carlo trials of each mechanism.

**S1 Video. M2 Maximum UL-Similarity.** This video demonstrates the run for which we obtained maximum UL-Similarity for Mechanism 2. The maximum UL-Similarity occurs around 6 seconds into the video. Gray and burgundy fluctuate toward the end, and the northwest corner of the region undergoes immediate change, unlike in Mechanisms 3 and 4.

**S2 Video. M3 Maximum UL-Similarity.** Here, we show the run for maximum UL-Similarity for Mechanism 3. Blue*, gray*, and burgundy* were triggered around 13 seconds, when a blue TU reached the northernmost row of active TUs. Maximum UL-Similarity was achieved at around 16 seconds.

**S3 Video. M4 Maximum UL-Similarity.** We provide the run for maximum UL-Similarity for Mechanism 4. The trigger is noticeable, as in S2 Video. Maximum UL-Similarity was again achieved at around 16 seconds.

**S4 Video. M4 Maximum Similarity.** This video shows the run where maximum Similarity was achieved for Mechanism 4. The trigger is noticeable, as in S2 and S3 Videos. Maximum Similarity occurs around the 18-second mark.

**S5 Video. M4 Sample Video.** This video is smooth and appears biomimetic- it represents one of the better Mechanism 4 runs. Maximum UL-Similarity occurs around 17 seconds into the video.

**S6 Video. M4 Sample Video 2.** We show another example of a Mechanism 4 run. Here, maximum UL- Similarity is achieved a bit earlier, around the 12-second mark.

## References

[1] Einhorn TA. Enhancement of fracture-healing. J Bone Joint Surg Am. 1995 Jun;77(6): 940–956. http://journals.lww.com/jbjsjournal/Citation/1995/06000/Enhancement_of_fracture_healing_.16.aspx

[2] Harvey N, Dennison E, Cooper C. Osteoporosis: impact on health and economics. Nat Rev Rheumatol. 2010 Feb;6(2): 99–105. https://www.nature.com/nrrheum/journal/v6/n2/full/nrrheum.2009.260.html

[3] Nandi SK, Roy S, Mukherjee P, Kundu B, De DK, Basu D. Orthopaedic applications of bone graft & graft substitutes: a review. Indian J Med Res, 2010 Jul;132: 15–30. http://imsear.li.mahidol.ac.th/handle/123456789/135534

[4] Hustedt JW, Blizzard DJ. The controversy surrounding bone morphogenetic proteins in the spine: a review of current research. Yale J Biol Med. 2014 Dec 12;87(4): 549–561. https://www.ncbi.nlm.nih.gov/pmc/articles/PMC4257039/

[5] Turner WG. The Use of the Bone Graft in Surgery. Can Med Assoc J. 1915 Feb;5(2): 103–109. https://www.ncbi.nlm.nih.gov/pmc/articles/PMC1487122/

[6] Vodovotz Y, An G. Chapter 1.1. Interesting Times: The Translational Dilemma and the Need for Translational Systems Biology of Inflammation. In: Vodovotz Y, An G, authors. Translational Systems Biology. London: Academic Press; 2014. pp. 3–8. https://www.elsevier.com/books/translational-systems-biology/vodovotz/978-0-12-397884-4

[7] Ioannidis JP. To replicate or not to replicate: the case of pharmacogenetic studies: Have pharmacogenomics failed, or do they just need larger-scale evidence and more replication? Circ Cardiovasc Genet. 2013 Aug;6(4):413–418. http://circgenetics.ahajournals.org/content/6/4/413

[8] Begley CG, Ellis LM. Drug development: Raise standards for preclinical cancer research. Nature. 2012 Mar 28;483(7391): 531–533. https://www.nature.com/nature/journal/v483/n7391/full/483531a.html

[9] Einhorn TA, Gerstenfeld LC. Fracture healing: mechanisms and interventions. Nat Rev Rheumatol. 2015 Jan 1;11(1): 45–54. http://www.nature.com/nrrheum/journal/v11/n1/abs/nrrheum.2014.164.html

[10] Bais M, McLean J, Sebastiani P, Young M, Wigner N, Smith T, Kotton DN, Einhorn TA, Gerstenfeld LC. Transcriptional Analysis of Fracture Healing and the Induction of Embryonic Stem Cell–Related Genes. PLoS One. 2009 May 5;4(5): e5393. http://journals.plos.org/plosone/article?id=10.1371/journal.pone.0005393

[11] Pezzulo G, Levin M. Top-down models in biology: explanation and control of complex living systems above the molecular level. J R Soc Interface. 2016 Nov 1;13(124): 20160555. http://rsif.royalsocietypublishing.org/content/13/124/20160555

[12] Fonstad MA. Cellular automata as analysis and synthesis engines at the geomorphology–ecology interface. Geomorphology. 2006 Jul 30;77(3): 217–34. http://www.sciencedirect.com/science/article/pii/S0169555X06000134

[13] Anné J, Edwards NP, Wogelius RA, Tumarkin-Deratzian AR, Sellers WI, van Veelen A, Bergmann U, Sokaras D, Alonso-Mori R, Ignatyev K, Egerton VM. Synchrotron imaging reveals bone healing and remodelling strategies in extinct and extant vertebrates. J R Soc Interface. 2014 Jul 6;11(96): 20140277. http://rsif.royalsocietypublishing.org/content/11/96/20140277

[14] Marino S, Cilfone NA, Mattila JT, Linderman JJ, Flynn JL, Kirschner DE. Macrophage polarization drives granuloma outcome during Mycobacterium tuberculosis infection. Infection and immunity. 2015 Jan 1;83(1):324–38. http://iai.asm.org/content/83/1/324

[15] Gardiner BS, Wong KK, Joldes GR, Rich AJ, Tan CW, Burgess AW, Smith DW. Discrete element framework for modelling extracellular matrix, deformable cells and subcellular components. PLoS Comp Biol. 2015 Oct 9;11(10):e1004544. http://journals.plos.org/ploscompbiol/article?id=10.1371/journal.pcbi.1004544

[16] Ziraldo C, Solovyev A, Allegretti A, Krishnan S, Henzel MK, Sowa GA, Brienza D, An G, Mi Q, Vodovotz Y. A computational, tissue-realistic model of pressure ulcer formation in individuals with spinal cord injury. PLoS Comp Biol. 2015 Jun 25;11(6):e1004309. http://journals.plos.org/ploscompbiol/article?id=10.1371/journal.pcbi.1004309

[17] Slade Shantz JA, Yu YY, Andres W, Miclau T 3rd, Marcucio R. Modulation of macrophage activity during fracture repair has differential effects in young adult and elderly mice. J Orthop Trauma. 2014;28 Suppl 1: S10–4. http://europepmc.org/abstract/MED/24378434

[18] Jiao H, Xiao E, Graves DT. Diabetes and Its Effect on Bone and Fracture Healing. Curr Osteoporos Rep. 2015 Oct;13(5): 327–335. https://link.springer.com/article/10.1007%2Fs11914-015-0286-8

[19] Scolaro JA, Schenker ML, Yannascoli S, Baldwin K, Mehta S, Ahn J. Cigarette smoking increases complications following fracture: a systematic review. J Bone Joint Surg Am. 2014 Apr 16;96(8): 674–681. https://insights.ovid.com/pubmed?pmid=24740664

[20] Patton CM, Powell AP, Patel AA. Vitamin D in orthopaedics. J Am Acad Orthop Surg. 2012 Mar;20(3): 123–129. https://insights.ovid.com/pubmed?pmid=22382284

[21] Hunt CA, Ropella GE, Lam TN, Tang J, Kim SH, Engelberg JA, Sheikh-Bahaei S. At the biological modeling and simulation frontier. Pharm Research. 2009 Nov 1;26(11): 2369–400. https://link.springer.com/article/10.1007/s11095-009-9958-3

[22] Tang J, Hunt CA. Identifying the rules of engagement enabling leukocyte rolling, activation, and adhesion. PLoS Comp Biol. 2010 Feb 19;6(2):e1000681. http://journals.plos.org/ploscompbiol/article?id=10.1371/journal.pcbi.1000681

[23] Lam TN, Hunt CA. Mechanistic insight from in silico pharmacokinetic experiments: roles of P-glycoprotein, Cyp3A4 enzymes, and microenvironments. J Pharmacol Exp Therap. 2010 Feb 1;332(2): 398–412. http://jpet.aspetjournals.org/content/332/2/398

[24] Sheikh-Bahaei S, Hunt CA. Enabling clearance predictions to emerge from in silico actions of quasi-autonomous hepatocyte components. Drug Metab Dispos. 2011 Oct 1;39(10): 1910–20. http://dmd.aspetjournals.org/content/39/10/1910

[25] Kim SHJ, Jackson AJ, Hunt CA (2014) In silico, experimental, mechanistic model for extended-release Felodipine disposition exhibiting complex absorption and a highly variable food interaction. PLoS ONE Sept 30;9(9): e108392. http://journals.plos.org/plosone/article?id=10.1371/journal.pone.0108392

[26] Petersen BK, Ropella GE, Hunt CA. Toward modular biological models: defining analog modules based on referent physiological mechanisms. BMC Sys Biol. 2014 Aug 16;8(1):95. https://bmcsystbiol.biomedcentral.com/articles/10.1186/s12918-014-0095-1

[27] Smith AK, Petersen BK, Ropella GE, Kennedy RC, Kaplowitz N, Ookhtens M, Hunt CA. Competing Mechanistic Hypotheses of Acetaminophen-Induced Hepatotoxicity Challenged by Virtual Experiments. PLoS Comput Biol. 2016 Dec 16;12(12): e1005253. http://journals.plos.org/ploscompbiol/article?id=10.1371/journal.pcbi.1005253

[28] Luke S, Cioffi-Revilla C, Panait L, Sullivan K, Balan G. MASON: A Multiagent Simulation Environment. Simulation. 2005;81(7): 517–527. http://dl.acm.org/citation.cfm?id=1086877

[29] Stich T, Linz C, Wallraven C, Cunningham D, Magnor M. Perception-motivated interpolation of image sequences. ACM Trans Appl Percept. 2011 Jan 1;8(2): 11. http://dl.acm.org/citation.cfm?id=1870079

[30] Petersen BK, Ropella GE, Hunt CA. Virtual experiments enable exploring and challenging explanatory mechanisms of immune-mediated P450 down-regulation. PLoS ONE. 2016;11: e0155855. http://journals.plos.org/plosone/article?id=10.1371/journal.pone.0155855

[31] Hunt CA, Kennedy RC, Kim SH, Ropella GE. Agent-based modeling: a systematic assessment of use cases and requirements for enhancing pharmaceutical research and development productivity. Wiley Interdiscip Rev Syst Biol Med. 2013 Jul 1;5(4): 461–80. http://onlinelibrary.wiley.com/doi/10.1002/wsbm.1222/full

[32] Kirschner DE, Hunt CA, Marino S, Fallahi-Sichani M, Linderman JJ. Tuneable resolution as a systems biology approach for multi-scale, multi-compartment computational models. Wiley Interdiscip Rev Syst Biol Med. 2014;6: 289–309. http://onlinelibrary.wiley.com/doi/10.1002/wsbm.1270/abstract

[33] Bartha P. Analogy and Analogical Reasoning. The Stanford Encyclopedia of Philosophy (Fall 2013 Edition). 2013. Available from: http://plato.stanford.edu/archives/fall2013/entries/reasoning-analogy/

[34] Frigg R, Hartmann, S. Models in Science. The Stanford Encyclopedia of Philosophy (Fall 2012 Edition). 2012. Available from: http://plato.stanford.edu/archives/fall2012/entries/models-science/

[35] Callus Analog Framework. Available: https://simtk.org/projects/callusanalog

[36] Hinton RJ, Jing Y, Jing J, Feng JQ. Roles of Chondrocytes in Endochondral Bone Formation and Fracture Repair. J Dent Res. 2017 Jan;96(1): 23–30. http://journals.sagepub.com/doi/abs/10.1177/0022034516668321

[37] Wong SA, Hu D, Miclau T, Bahney C, Marcucio R. Trans differentiation of Chondrocytes to Osteoblasts during Endochondral Ossification in the Healing Mandible. FASEB J. 2016 Apr;30(1). http://www.fasebj.org/content/30/1_Supplement/1039.11

[38] Shapiro IM, Adams CS, Freeman T, Srinivas V. Fate of the hypertrophic chondrocyte: microenvironmental perspectives on apoptosis and survival in the epiphyseal growth plate. Birth Defects Res C Embryo Today. 2005 Dec;75(4): 330–9. http://onlinelibrary.wiley.com/doi/10.1002/bdrc.20057/abstract

[39] Hu DP, Ferro F, Yang F, Taylor AJ, Chang W, Miclau T, Marcucio RS, Bahney CS. Cartilage to bone transformation during fracture healing is coordinated by the invading vasculature and induction of the core pluripotency genes. Development. 2017 Jan 15;144(2): 221–234. http://dev.biologists.org/content/144/2/221

[40] Hunt CA, Ropella GE, ning Lam T, Gewitz AD. Relational grounding facilitates development of scientifically useful multiscale models. Theor Biol Med Mod. 2011 Sep 27;8(1):35. https://tbiomed.biomedcentral.com/articles/10.1186/1742-4682-8-35

[41] Assembla. Available: https://www.assembla.com

[42] Walpole J, Papin JA, Peirce SM. Multiscale computational models of complex biological systems. Annu Rev Biomed Eng. 2013;15: 137–54. http://www.annualreviews.org/doi/abs/10.1146/annurev-bioeng-071811-150104

[43] Hoff P, Gaber T, Strehl C, Schmidt-Bleek K, Lang A, Huscher D, Burmester GR, Schmidmaier G, Perka C, Duda GN, Buttgereit F. Immunological characterization of the early human fracture hematoma. Immunol Res. 2016 Dec;64(5-6): 1195–1206. https://link.springer.com/article/10.1007%2Fs12026-016-8868-9

[44] Gómez-Picos P, Eames BF. On the evolutionary relationship between chondrocytes and osteoblasts. Front Genet. 2015 Sep 23;6: 297. http://journal.frontiersin.org/article/10.3389/fgene.2015.00297/full

[45] Miclau KR, Brazina SA, Bahney CS, Hankenson KD, Hunt TK, Marcucio RS, Miclau T. Stimulating Fracture Healing in Ischemic Environments: Does Oxygen Direct Stem Cell Fate during Fracture Healing? Front Cell Dev Biol. 2017 May 4;5: 45. http://journal.frontiersin.org/article/10.3389/fcell.2017.00045/full

[46] OReilly A, Hankenson KD, Kelly DJ. A computational model to explore the role of angiogenic impairment on endochondral ossification during fracture healing. Biomech Model Mechanobiol. 2016 Oct;15(5): 1279–94. https://link.springer.com/article/10.1007%2Fs10237-016-0759-4

[47] Kaae S, Johansen H. Does simple mastectomy followed by irradiation offer survival comparable to radical procedures? Int J Radiat Oncol Biol Phys. 1977 Nov-Dec;2(11-12): 1163–6. https://linkinghub.elsevier.com/retrieve/pii/0360-3016(77)90126-2

[48] Colnot C, Thompson Z, Miclau T, Werb Z, Helms JA. Altered fracture repair in the absence of MMP9. Development. 2003 Sep;130(17): 4123–4133. http://dev.biologists.org/content/130/17/4123.long

[49] Gerber HP, Vu TH, Ryan AM, Kowalski J, Werb Z, Ferrara N. VEGF couples hypertrophic cartilage remodeling, ossification and angiogenesis during endochondral bone formation. Nat Med. 1999 Jun;5(6): 623–628. https://www.nature.com/nm/journal/v5/n6/full/nm0699_623.html

[50] Colnot CI, Helms JA. A molecular analysis of matrix remodeling and angiogenesis during long bone development. Mech Dev. 2001 Feb;100(2): 245–250. http://www.sciencedirect.com/science/article/pii/S0925477300005323?via%3Dihub

[51] Maes F, Vandermeulen D, Suetens P. Medical image registration using mutual information. Proc of IEEE. 2003 Oct; 91(10): 1699–1722. http://ieeexplore.ieee.org/abstract/document/1232201/

